# Transcription factor-based transdifferentiation of human embryonic to trophoblast stem cells

**DOI:** 10.1101/2021.08.18.456785

**Authors:** Paula A. Balestrini, Abdelbaki Ahmed, McCarthy Afshan, Liani Devito, Claire E. Senner, Alice E. Chen, Prabhakaran Munusamy, Paul Blakeley, Kay Elder, Phil Snell, Leila Christie, Paul Serhal, Rabi A. Odia, Mahesh Sangrithi, Kathy K. Niakan, Norah M.E. Fogarty

## Abstract

During the first week of development, human embryos form a blastocyst comprised of an inner cell mass and trophectoderm (TE) cells, the latter of which are progenitors of placental trophoblast. Here we investigated the expression of transcripts in the human TE from early to late blastocyst stages. We identified enrichment of transcription factors *GATA2*, *GATA3*, *TFAP2C* and *KLF5* and characterised their protein expression dynamics across TE development. By inducible overexpression and mRNA transfection we determined that these factors, together with MYC, are sufficient to establish induced trophoblast stem cells (iTSCs) from primed human embryonic stem cells. These iTSCs self-renew and recapitulate morphological characteristics, gene expression profiles, and directed differentiation potential similar to existing human TSCs. Systematic omission of each, or combinations of factors, revealed the critical importance of GATA2 and GATA3 for iTSC transdifferentiation. Altogether, these findings provide insights into the transcription factor network that may be operational in the human TE and broaden the methods for establishing cellular models of early human placental progenitor cells, which may be useful in the future to model placental-associated diseases.

**Summary statement:** Transcriptional analysis of human blastocysts reveals transcription factors sufficient to derive induced trophoblast stem cells from primed human embryonic stem cells.

## Introduction

The correct functioning of the human placenta is crucial to ensure a healthy pregnancy outcome. However, despite this critical role, it is one of the least understood organs. After fertilisation the zygote undergoes a series of cleavage cell divisions to form a tight ball of cells called a morula. The outer cells of the morula become polarised which leads to inactivation of the Hippo signalling pathway and allows the translocation of YAP1 into the nucleus where, together with TEAD4, it drives transcriptional activation of the trophectoderm (TE) programme (Gerri et al., 2020). The human blastocyst implants into the uterine lining approximately 7-9 days after fertilisation. The TE generates the first trophoblast lineages: the mononuclear cytotrophoblast cells (CTB) and the multinucleated primitive syncytiotrophoblast (STB) that contribute to the fetal portion of the placenta. The primitive STB is the initial invading interface and enzymatically degrades the uterine lining allowing for implantation of the conceptus (Hertig et al., 1956). CTB cells are the precursor for trophoblast cell types in the placenta. CTB cells sustain the syncytiotrophoblast across gestation by dividing asymmetrically, with one daughter cell fusing with the overlying syncytiotrophoblast and the other daughter remaining in the proliferative pool (Baczyk et al., 2006). The syncytiotrophoblast performs key functions including providing a physical and immunological barrier to pathogens, transporting nutrients and gaseous exchange, as well as producing and secreting hormones required to adapt the maternal physiology to the pregnancy (Bansal et al., 2012; Costa, 2016). CTB cells are also the precursor of extravillous trophoblast cells (EVTs) which invade the decidua to interact with maternal immune cells and remodel spiral arteries to establish blood supply to the placenta (Pijnenborg et al., 1980).

Human trophoblast stem cells (hTSCs) derived from blastocysts and first trimester placental villous tissue provide a paradigm for elucidating molecular mechanisms regulating trophoblast development (Okae et al., 2018). hTSCs show expression of well-established markers of trophoblast including TP63 (Okae et al., 2018), GATA3 (Chiu and Chen, 2016) and TEAD4 (Soncin et al., 2018) and can be directed to differentiate into both EVT and STB, as identified by the expression of markers HLA-G (Apps et al., 2008) and SDC1 (Jokimaa et al., 1998), respectively. In addition, the establishment of a trophoblast organoid culture system has allowed hTSCs to be cultured in 3D, somewhat resembling the architecture of villous CTB cells and STB of the placenta (Haider et al., 2018; Turco et al., 2018). Most recently, induced trophoblast stem cells (iTSCs) which resemble primary tissue-derived hTSCs have been captured in the process of transdifferentiation of fibroblasts to human embryonic stem cells (hESCs) in naïve cell culture conditions (Castel et al., 2020; Cinkornpumin et al., 2020; Dong et al., 2020; Liu et al., 2020; Naama et al., 2023). In addition, established naïve ES cells cultured cultured in conditions which inhibit ERK and Nodal signalling also yield iTSCs (Guo et al., 2021). These *in vitro* models can be utilised in studies of trophoblast biology and have the potential to further inform our understanding of molecular mechanisms, transcription factors and signalling pathways regulating human trophoblast biology.

Studies of mouse preimplantation embryogenesis have elucidated mechanisms regulating lineage specification and maintenance of the TE in this species (Hemberger et al., 2020). In turn, this knowledge has led to the development of strategies to capture their *in vitro* counterparts. Blastocyst-derived TSCs can be isolated from the polar TE of blastocyst outgrowths or post-implantation extraembryonic ectoderm cultured in TSC media supplemented with FGF4 and heparin which recapitulates the signalling environment of the *in vivo* TSC compartment at the post-implantation stage (Tanaka et al., 1998). Identification of key transcription factors regulating mouse TE lineage specification and maintenance has also informed transdifferentiation strategies. Single overexpression of *Cdx2* or *Eomes* in TSC media is sufficient to induce differentiation of mouse ESCs to trophoblast-like cells (Niwa et al., 2005). Transient ectopic expression of the combination of TE-associated transcription factors *Gata3*, *Eomes*, *Tcfap2c*, with either *Ets2* or *Myc*, has been shown to reprogramme fibroblasts into mouse iTSCs (Benchetrit et al., 2015; Kubaczka et al., 2015). These alternatively derived cells recapitulate the morphology, epigenomes and gene expression patterns of blastocyst-derived TSCs and contribute exclusively to the placenta following blastocyst injection.

Here we perform a time-course RNA-sequencing analysis of human TE development and explored gene expression profiles of the human preimplantation TE to identify associated transcription factors that may be employed as transdifferentiation factors to convert primed hESCs to iTSCs. We identified four transcription factors (*GATA2*, *GATA3*, *TFAP2C* and *KLF5*) that are enriched in the TE lineage of human blastocysts compared with EPI and PE. We confirmed high expression of these factors in TE cells at all stages analysed at both the transcript and the protein level. Similar to strategies used in the mouse, we hypothesized that the transient ectopic expression of these transcription factors, along with the reprogramming enhancing factor *MYC*, would facilitate the establishment of human iTSCs. Here, we demonstrate using both a doxycycline-inducible approach and modified mRNA transfection that the overexpression of these factors promotes transdifferentiation of hESCs from primed condition to iTSCs. The iTSCs derived can be maintained stably in culture and recapitulate the expression of markers that have been used previously to identify embryo and primary placental-derived hTSCs and trophoblast (Lee et al., 2016; Okae et al., 2018). RNA-seq analysis reveals that iTSCs are transcriptionally similar to existing hTSCs and primary CTB cells. When subjected to directed differentiation protocols iTSCs can both undergo syncytialisation to form multinucleated structures that secrete a syncytiotrophoblast-associated peptide hormone, human chorionic gonadotropin (hCG) as well as forming elongated extravillous trophoblast cells (EVTs). Altogether this reveals a novel strategy for the establishment of iTSCs and significantly expands the repertoire of cells to model placenta biology *in vitro*.

## Results

### Identification and validation of candidate transcription factors

In this study we sought to devise a transdifferentiation strategy to generate iTSCs informed by the expression of transcription factors that are enriched in the TE. We initially sought to determine which genes overlap in their expression in the TE across preimplantation stages of human development, reasoning that commonly expressed transcripts may be required for the establishment and maintenance of the TE and thereby may facilitate iTSC transdifferentiation. We cultured human embryos from embryonic day 5 when the blastocyst is first established and TE cells are discernible, until embryonic day 7 which is the latest stage we can culture preimplantation blastocysts to *in vitro* (Fig. 1A). We performed RNA-sequencing analysis to determine the gene expression profile for each developmental stage (Table S1). Bulk RNA-sequencing analysis allowed us to identify lowly expressed genes, including transcription factors, which often drop out from single cell RNA-sequencing data (Kharchenko et al., 2014), and had the additional benefit of minimising the numbers of embryos used in this study. We did not observe expression of molecular markers associated with the epiblast or primitive endoderm, such as *NANOG* or *SOX17*, in TE samples indicating we did not have contamination from ICM cells (Table S1). t-Distributed stochastic neighbour embedding (t-SNE) dimensionality reduction analysis indicated that plotting the first two dimensions of the t-SNE separates TE samples into three groups corresponding to the developmental time-points analysed (Fig. 1B). This was confirmed by a principal component analysis (PCA) that showed that when the first five principal components (PC) are plotted against each other, PC1 separates TE samples into the three developmental time-points (Fig. S1A). We compared the global gene expression patterns at these stages of development to identify the transcripts that are commonly expressed and those unique to each stage (Table S2). We found that most transcripts, 4729 genes, were expressed in common at the stages analysed compared with 576, 387, and 250 genes uniquely expressed at day 5, 6, and 7, respectively (Fig. 1C). This suggests that once initiated the TE programme remains transcriptionally consistent across preimplantation development. Functional enrichment analysis using GeneOntology (Ashburner et al., 2000; Gene Ontology, 2021; Mi et al., 2019) and REACTOME databases (Jassal et al., 2020) revealed enrichment in metabolic processes, translational activities and molecule binding which is consistent with the rapid expansion of the TE (Fig. 1D). Upon further inspection of the gene lists associated with these terms we observed an enrichment in genes that are required for efficient oxidative phosphorylation, including genes encoding complexes of the electron transport chain (*MT-ND1*, *SDHD*, *CYC1*, *UQCRQ, UQCRC1*, *ATP 5*,*EATP 5* a*I* nd *ATP5L*). In addition, we observed an enrichment of genes involved in mitochondrial morphology. Interestingly while we observed enrichment of genes regulating mitochondrial fusion (*MFN1*, *M F N 1*and *OPA1*), we did not detect expression of genes regulating mitochondrial fission (*DRP1*). In all, this suggests that mitochondrial fusion in the TE may be essential for the electron transport chain favouring oxidative phosphorylation. This is consistent with the observation that OXPHOS activity increases in developing mammalian blastocysts and correlates with the capacity of the embryo to develop to term following implantation (Gardner et al., 1998; Houghton, 2006; Leese et al., 2008). The genes expressed in common included *PPARG* which is expressed in CTBs and regulates differentiation (Murthi et al., 2013), a marker of CTBs *SPINT1 (Kohama et al., 2012)* and *TCF7L1,* a key transcription factor of the WNT signalling pathway that is implicated in trophoblast proliferation (Meinhardt et al., 2014) (Fig. 1E). *ACE2*, *APOE* and the receptor protein-tyrosine kinase *EFNA4* were detected at the earlier time-points analysed whereas *WLS,* a core component of the WNT secretion pathway *(Yu et al., 2014), VHL* a regulator of placenta vasculogenesis (Genbacev et al., 2001) and *HAND1* a transcription factor involved in branching morphogenesis during mouse placentation (Riley et al., 1998) were enriched at later timepoints.

**Figure 1.**
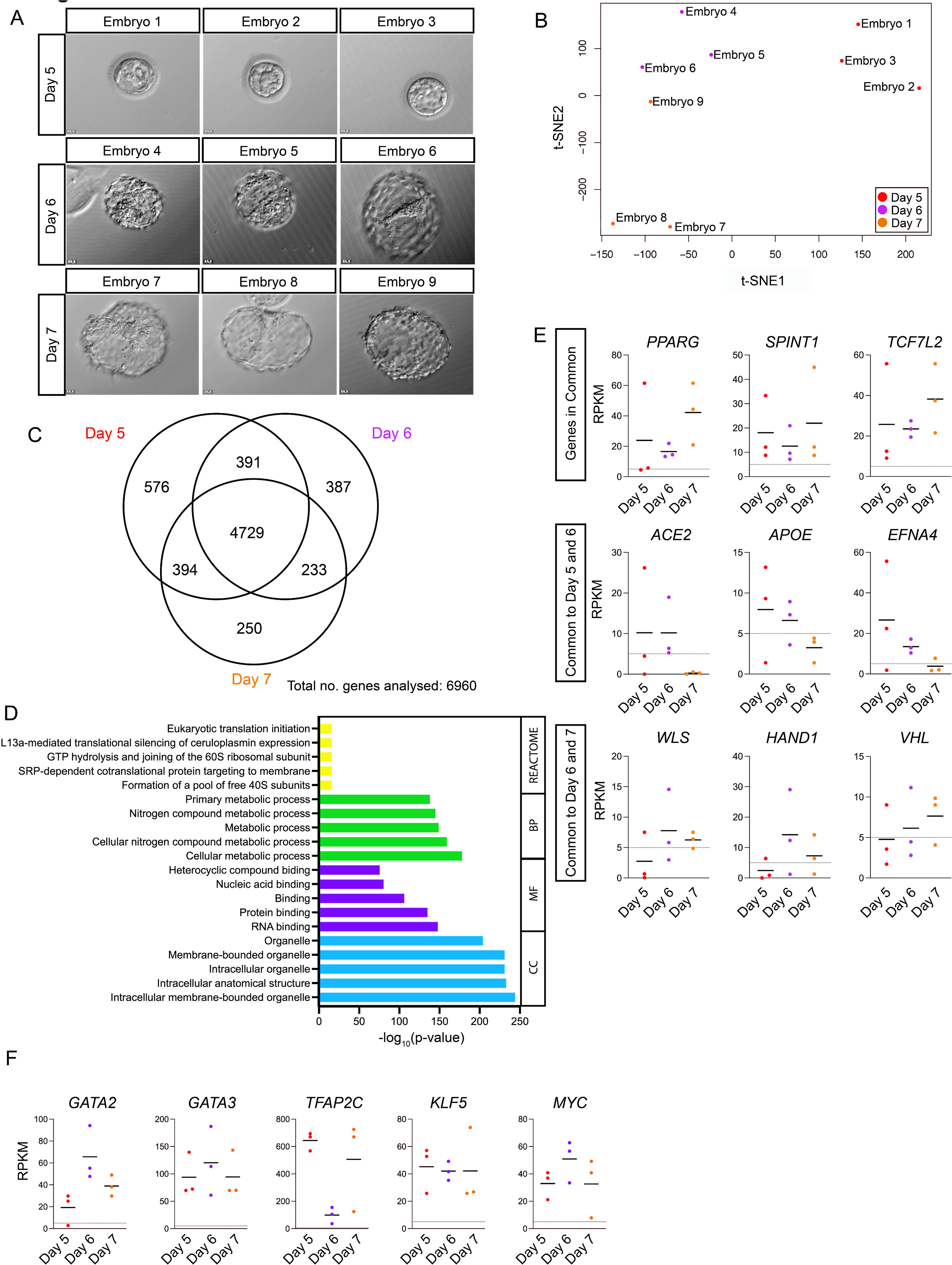
Transcriptome analysis of TE in human blastocysts. (A) Brightfield images of human blastocysts microdissected for TE isolation ahead of RNA-sequencing analysis. (B) t-Distributed Stochastic Neighbour Embedding (t-SNE) plot based on VST-normalisation of the top 6000 most variable expressed genes. A colour coded sample key is provided. (C) Venn diagram indicating the number of transcripts with overlap or unique expression at day 5, day 6 and day 7 in the mural TE (RPKM>5). (D) The most enriched GO terms for genes associated with biological process (BP), cellular component (CC), molecular function (MF), and REACTOME pathways associated with genes expressed in common are shown. (E) Scatterplots showing selected genes that are expressed in common at all stages analysed, expressed in common at days 5 and 6, or expressed in common at days 6 and 7. Horizontal line denotes the mean RPKM value of each timepoint, dotted line shows RPKM value = 5 (F) Scatterplots showing expression of *GATA2*, *GATA3*, *TFAP2C*, *KLF5* and *MYC* at days 6 and 7. Horizontal line denotes the mean RPKM value of each timepoint, dotted line shows RPKM value = 5.

We collected a comprehensive transcription factor annotation from Human Transcription Factor Database (Hu et al., 2019) and cross-referenced this against the list of transcripts detected in common at all stages analysed. This identified 232 transcription factors expressed in the TE (Table S3). We similarly mined our previously published single-cell RNA-seq datasets to identify transcription factors that were significantly enriched in the TE compared to the EPI and PE cells(Blakeley et al., 2015). We further refined these lists of TE-associated transcription factors by screening for transcription factors with a known role in mouse TE development (Kuckenberg et al., 2010; Ralston et al., 2010) or a suggested role in the human placenta. In all, from these analyses we identified four transcription factors for further investigation as candidate transdifferentiation factors: *GATA2, GATA3, TFAP2C* and *KLF5* (Fig. 1F) (Table S4).

From our analysis of single-cell RNA-seq data using our previously published Shiny App programme(Blakeley et al., 2015; Wamaitha et al., 2020), we found that *GATA2*, *GATA3* and *TFAP2C* share a common pattern of expression, with an onset of embryonic transcription shortly prior to TE initiation at the 8-cell stage (Gerri et al., 2020) and high expression maintained as development proceeds. By contrast, *KLF5* is expressed in human zygotes and its expression increases from the 8-cell stage. Analysis of lineage-specific gene expression patterns showed that *GATA2* and *GATA3* are enriched specifically in the TE. While *TFAP2C* is detected in the TE, it is also detected in the EPI, which we previously confirmed at the protein level. *KLF5* is detected in all three lineages, with the most abundant expression in the TE (Blakeley et al., 2015).

We next performed immunofluorescence analysis of these transcription factors in human embryos cultured from embryonic day 5 to 7. At all stages analysed, GATA2 protein expression was detected in the nuclei in TE cells, which were identified by both their position within the blastocyst and the absence of NANOG expression (Fig. 2A). Similarly, GATA3 protein was detected in TE cells across the stages analysed (Fig. 2B). The pattern of GATA2 and GATA3 expression is similar to what has been reported in mouse embryos (Home et al., 2017) and is consistent with our previous analysis of GATA3 expression (Gerri et al., 2020). At all stages analysed nuclear KLF5 was detected in the TE, whereas the inner cell mass (ICM) cells showed some cytoplasmic expression (Fig. 2C). This pattern is similar to what has been described in the mouse, where KLF5 is expressed in all cells of preimplantation embryos with lower expression in ICM compared to the TE (Lin et al., 2010). By contrast, TFAP2C was detected at similar levels in both TE cells and NANOG-positive epiblast cells throughout all the blastocyst stages analysed. As we, and others, reported previously this contrasts with the expression pattern in the mouse where the homologue TCFAP2C is exclusively expressed in the TE (Fig. 2D) (Blakeley et al., 2015; Kuckenberg et al., 2010). Altogether, the enriched expression of GATA2, GATA3, TFAP2C and KLF5 in the TE suggests that these transcription factors may have a functional role within the TE transcriptional network.

**Figure 2.**
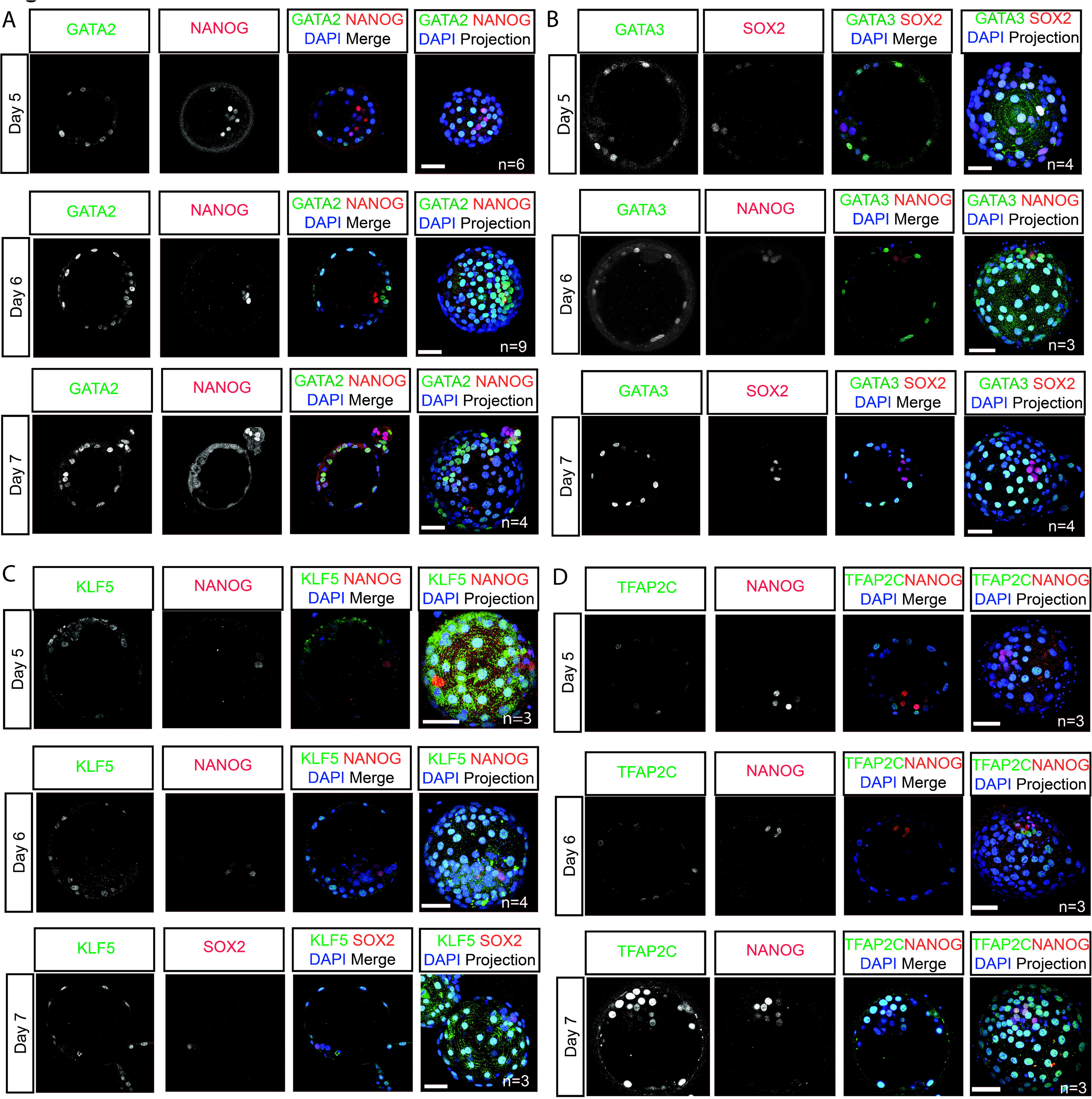
Identification of TE-associated transcription factors in the human blastocyst. Immunofluorescence analysis of (A) GATA2, (B) GATA3, (C) KLF5 and (D) TFAP2C (green) and DAPI nuclear staining (blue) in human blastocysts cultured to day 6 post-fertilisation. ICM cells are detected by either the expression of NANOG or SOX2 (red). Scale bars: 50 µm.

### Generation of 5F-iTSCs from hESCs using lentiviral doxycycline inducible system

We next evaluated whether the expression of these transcription factors was sufficient to facilitate the transdifferentiation of hESCs cultured in primed conditions directly to induced trophoblast stem cells (iTSCs) without a requirement to start from established naïve hESCs. We engineered doxycycline-inducible hESCs to overexpress *TFAP2C*, *KLF5*, *GATA3* and *GATA2* (Fig. 3A). We also induced the expression of *MYC* because we identified high expression of the transcript across the stages of TE development analysed (Table S4), and it has been shown to enhance the efficiency of cellular reprogramming in other contexts (Nakagawa et al., 2008). These cells are henceforth referred to as five factor-hESC (5F-hESCs).

**Figure 3.**
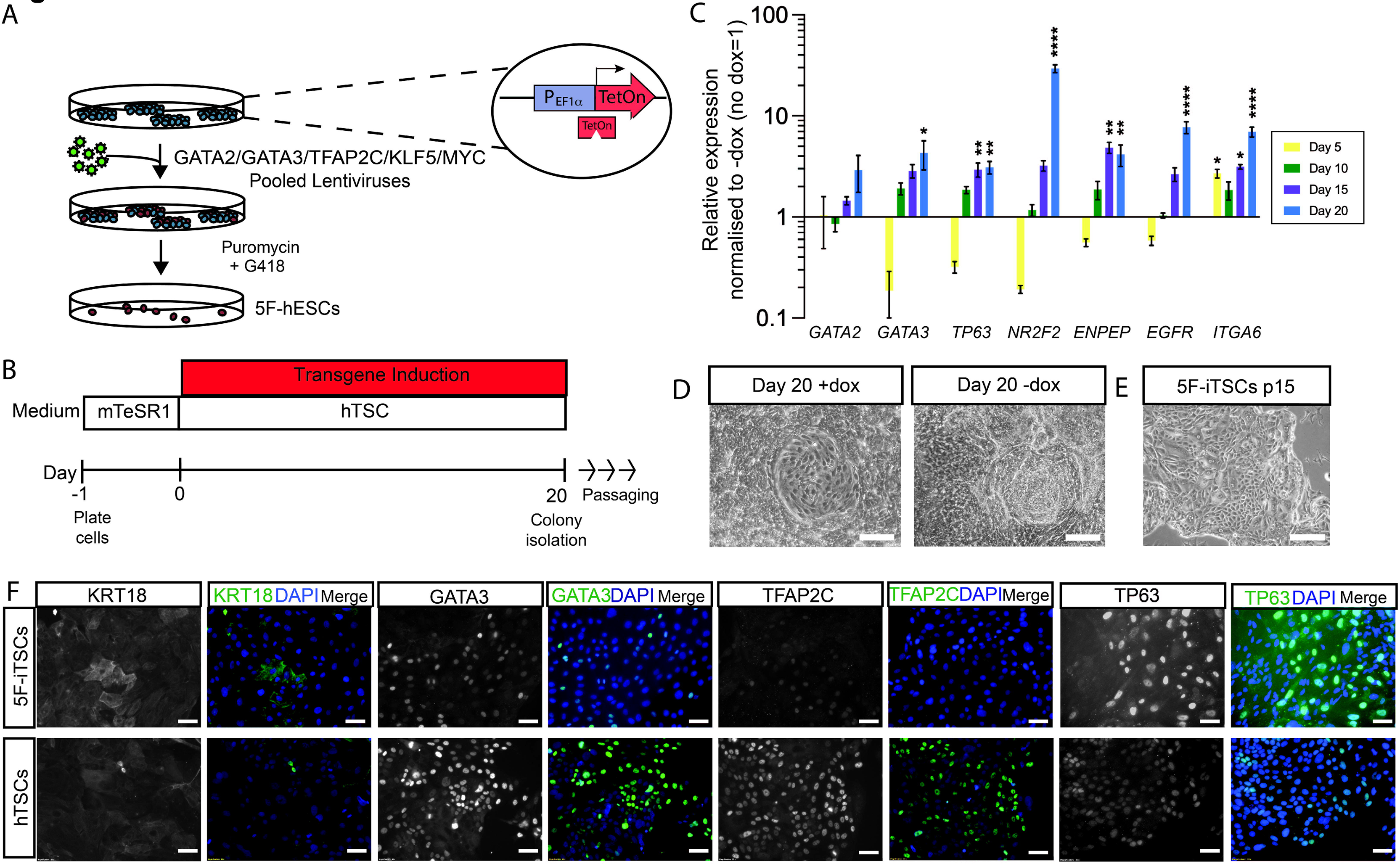
Induced of expression of GATA2, GATA3, TFAP2C and KLF5 for 20 days programs hESCs to hTSC-like cells. (A) Schematic diagram of the generation of GATA2, GATA3, TFAP2C, KLF5 and MYC inducible hESCs by lentiviral transduction. (B) Schematic representation of the strategy for transdifferentiation 5F-hESCs to hTSCs. 5F-hESCs were plated in mTeSR1 media for 24 h after which media was replaced with hTSC media. Doxycycline was administered daily for 20 days in hTSC media for transgene induction. (C) Time-course qRT-PCR analysis for the detection of endogenous *GATA2*, *GATA3*, *TP63*, *NR2F2*, *ENPEP*, *EGFR* and *ITGA6* in 5F-hESCs across 20 days of dox-induction in hTSC media. Relative expression is reflected as fold change over uninduced 5F-hESCs cultured in hTSC media normalized to *GAPDH*. Data are mean ± SEM of n=3 biological replicates analysed with a one-way ANOVA with Dunnett’s posthoc test (*p<0.05). (D) Brightfield images of 5F-hESCs cultured in hESCs cultured in hTSC media in the presence of dox for 20 days (+dox) and hTSC media alone for 20 days (-dox) (E) Brightfield images of stable 5F-iTSC lines derived from transgene overexpression grown for 15 passages. Scale bar: 200 µm. (F) Immunofluorescence analysis for the detection of KRT18, GATA3, TFAP2C and TP63 (green) and DAPI nuclear expression (blue) in stable 5F-iTSCs and in previously established control hTSCs. Scale bars: 50 µm).

To initiate the transdifferentiation process transgenes were induced by exposing 5F-hESCs to doxycycline in the presence of hTSC media for 20 days (Fig. 3B). This duration of transgene expression has been previously described for transcription factor-based transdifferentiation of murine fibroblasts to TSCs (Benchetrit et al., 2015). To gain insight into the effect of 5F induction during this time-period we performed a time-course RT-qPCR analysis for the detection of endogenous trophoblast-associated transcripts in induced 5F-hESCs compared with uninduced controls. *TP63* expression was upregulated after 10 days induction, contemporaneous with onset of endogenous *G A T A* e*3*xpression. *NR2F2* was subsequently upregulated (Fig. 3C). By day 20 we observed significant upregulation of the cell surface markers *ENPEP*, *ITGA6* and *EGFR* as well as further increases in expression levels *of GATA2, GATA3, TP63* and *NR2F2,* thus validating the duration of transgene induction for transdifferentiation. After 20 days, epithelial colonies with polygonal hTSC-like morphology were apparent within a heterogenous population as is similar to iPSC derivation and naïve programmming of hTSCs (Liu et al., 2020; Takahashi et al., 2007) (+dox, Fig. 3D). As a control, 5F-hESCs were cultured in hTSC media in the absence of doxycycline. In these conditions wide-spread cell proliferation and differentiation was observed (-dox, Fig. 3D). Colonies were manually selected and expanded upon in subsequent passages. Colonies continued to maintain their morphology and proliferate as iTSCs in the absence of doxycycline (Fig. 3E). Stable 5F-iTSC lines were generated and were maintained in culture for over 20 passages. By contrast, uninduced 5F-hESCs did not survive the first passage and underwent cell death. Immunofluorescence analysis of stable 5F-iTSCs indicated widespread expression of TE-associated keratin, KRT18, GATA3, TFAP2C and TP63 similar to exisiting hTSCs (Fig. 3E). By contrast, 5F-hESCs cultured in conventional primed conditions only showed some upregulation of KRT18 (Fig. S1C).

### Generation of 5F-iTSCs from hESCs using non-integrating modified mRNAs

Reprogramming via lentiviral transduction is reportedly an inefficient process (from 0.01% to 0.1%) (Wernig et al., 2008) and integrated constructs can be spontaneously silenced and reactivated during cell culture and differentiation(Ellis, 2005; Herbst et al., 2012). As an alternative approach to lentiviral transduction, in parallel we employed a strategy of transcription factor overexpression using chemically modified mRNAs, which has been used to generate transgene-free human iPSCs (Mandal and Rossi, 2013; Warren et al., 2010). Individual mRNAs encoding *GATA2, GATA3, TFAP2C, KLF5* and *MYC* were synthesised by *in vitro* transcription. Uridine and cytidine were substituted with the modified nucleotides pseudo-uridine and 5’methylcytidine to prevent cellular immune responses, and mRNA was capped with a modified 5’guanine cap to improve mRNA half-life. A cocktail of mRNA was made by pooling individual mRNAs in equal molar ratios and this was delivered into hESCs by lipofection every day for 20 days (Fig. 4A). Similar to what was observed with the doxycycline-inducible system, transfected cells underwent morphological change towards polygonal shaped cells within distinct colonies whereas mock transfected cells underwent widespread differentiation (Fig. 4B). After 20 days of transfection colonies were manually selected and stable iTSC lines were generated that could maintained in culture for over 20 passages (Fig. 4C). We next performed an in-depth analysis of mRNA generated 5F-iTSCs. qRT-PCR analysis of 5F-iTSCs confirmed significant upregulation of *GATA2*, *GATA3*, *EGFR, ENPEP,* and *TP63* compared with the starting population of hESCs (Fig. 4D). Immunofluorescence analysis of 5F-iTSCs confirmed widespread expression of trophoblast markers GATA2, GATA3, TFAP2C, TP63 and KRT18 (Fig. 4E). Previous studies have shown that strategies to derive TSCs from primed hESCs using growth factor modulation instead yields amnion-like cells (Guo et al., 2021; Io et al., 2021; Ohgushi et al., 2022). Here, we performed qRT-PCR analysis for the detection of aminon-associated transcripts *GABRP*, *HEY1*, *ISL1* and *WNT6.* While we detected low levels of expression of these transcripts, we found that levels of expression were equivalent to those detected in established hTSCs derived from blastocyst and placenta (Fig. 4F). qRT-PCR analysis of expression of the trophoblast-specific chromosome 19 microRNA cluster confirmed that 5F-iTSCs express miRNAs at levels comparable to established hTSCs (Fig. 4G). Methylation analysis of the *ELF5* promoter showed hypomethylation of the region in 5F-iTSCs and hTSCs, whereas the starting population of hESCs displayed hypermethylation (Fig. 4H). Altogether this suggests that the 5F-iTSCs exhibit key characteristics of *bona fide* hTSCs (Lee et al., 2016).

**Figure 4.**
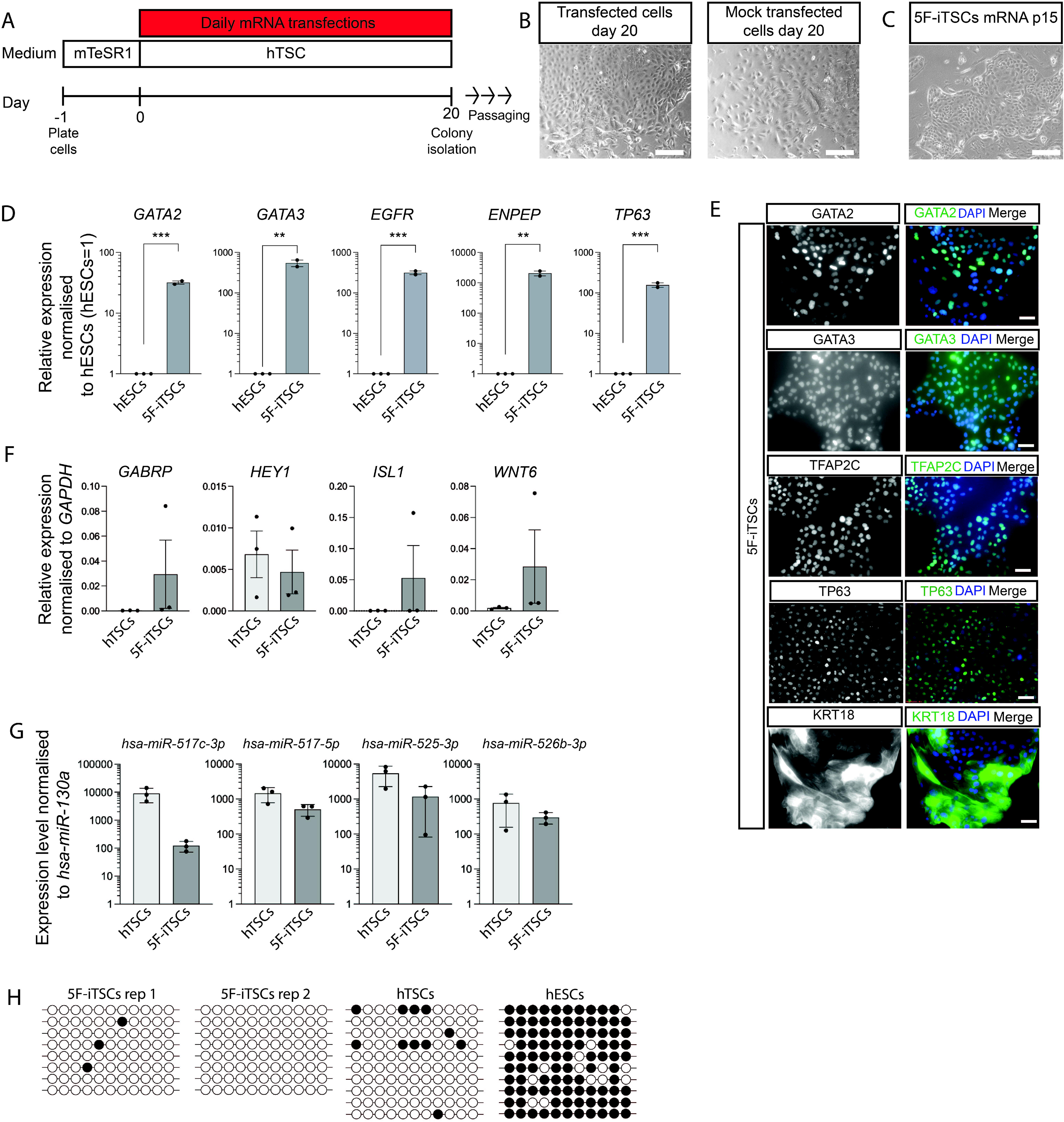
Characterisation of 5F-iTSCs programmed from primed hESCs using modified mRNAs. (A) Schematic representation of the strategy for transdifferentiation hESCs to iTSCs using modified mRNAs. hESCs were plated in mTeSR1 media for 24 h after which media was replaced with hTSC media. mRNAs encoding *GATA2*, *G A T A*, *T3FAP2C*, *K L F 5*and *M Y C* administered daily for 20 days via lipofection-mediated transfection. (B) Brightfield images of 5-factor mRNA-cocktail transfected hESCs (transfected) and mock transfected cells (mock) on day 20. Scale bar: 200 µm (C) Brightfield image of stable iTSCs derived from mRNA transdifferentiation grown for 15 passages. Scale bar: 200 µm. (D) RT-qPCR analysis for the detection of *GATA2*, *GATA3*, *ENPEP*, *EGFR* and *TP63* in stable 5F-iTSC. Relative expression is reflected as fold change over hESCs cultured in mTeSR media normalized to *GAPDH.* Data are mean ± SEM, n=2-3 biological replicates analysed with an unpaired one-tailed t-test (**p<0.01, ***p<0.005). (E) Immunofluorescence analysis for the detection of selected trophoblast markers GATA2, GATA3, TFAP2C, TP63 and KRT18 (green) and DAPI nuclear expression (blue) in stable 5F-iTSCs. Scale bar: 50 µm. (F) qRT-PCR for the detection of amnion-associated transcripts *GABRP*, *HEY1*, *ISL1* and *WNT6* in exisiting hTSCs and 5F-iTSCs. Relative expression is shown as ddCt values normalised to *GADPH*. Data are mean ± SEM, n=3 biological replicates analysed with an unpaired two-tailed t-test (n.s). (G) qRT-PCR for the detection of C19MC miRNAs in established hTSCs and 5F-iTSCs. Expression is normalised to *hsa-miR-130a*. Data are mean ± SEM, n=3 biological replicates analysed with an unpaired two-tailed t-test (not significant). (H) Bisulphite sequencing analysis of the *ELF5* promoter region. Filled circles indicate methylated cytosine residues. *ELF5* promoter is hypomethylated in 5F-iTSCs (n=2 biological replicates) and hTSCs, and methylated in hESCs.

### 5F-iTSCs are transcriptionally similar to previously established hTSCs and primary CTBs

We next compared global gene expression of mRNA-generated 5F-iTSCs to alternatively derived TSCs by integrating previously published datasets (Dong et al., 2020; Liu et al., 2020; Okae et al., 2018) as well as TE samples generated in this study, primary CTB cells (Haider et al., 2018) and primed hESCs (Dong et al., 2020). After adjusting for batch effects, we used the top 500 most variably expressed genes to perform dimensionality reduction analysis. Principal component analysis showed that plotting the first two principal components separated samples into three groups representing TE, hESCs and iTSCs together with all *in vitro* hTSCs and primary CTBs (Fig. 5A). To determine which samples were most similar with respect to genes with significant expression changes, we performed unsupervised hierarchical clustering. Consistent with the principal component analysis, three major clusters were observed: TE, hESCs, and iTSCs, *in vitro* hTSCs and primary CTBs (Fig 5B). Significantly, this analysis showed that the iTSCs generated in this study were most similar to hTSCs generated by Okae et al., followed by primary CTBs (Fig. 5B). Consistent with previous publications, we confirm that 5F-iTSCs and all hTSCs are transcriptionally similar to CTBs and distinct from the TE, suggesting that further refinement of hTSC medium is required to capture a developmentally earlier state. Together, these findings indicate that our 5F-iTSCs share a common transcriptomic profile with exisiting hTSCs and their in vivo cellular counterparts.

**Figure 5.**
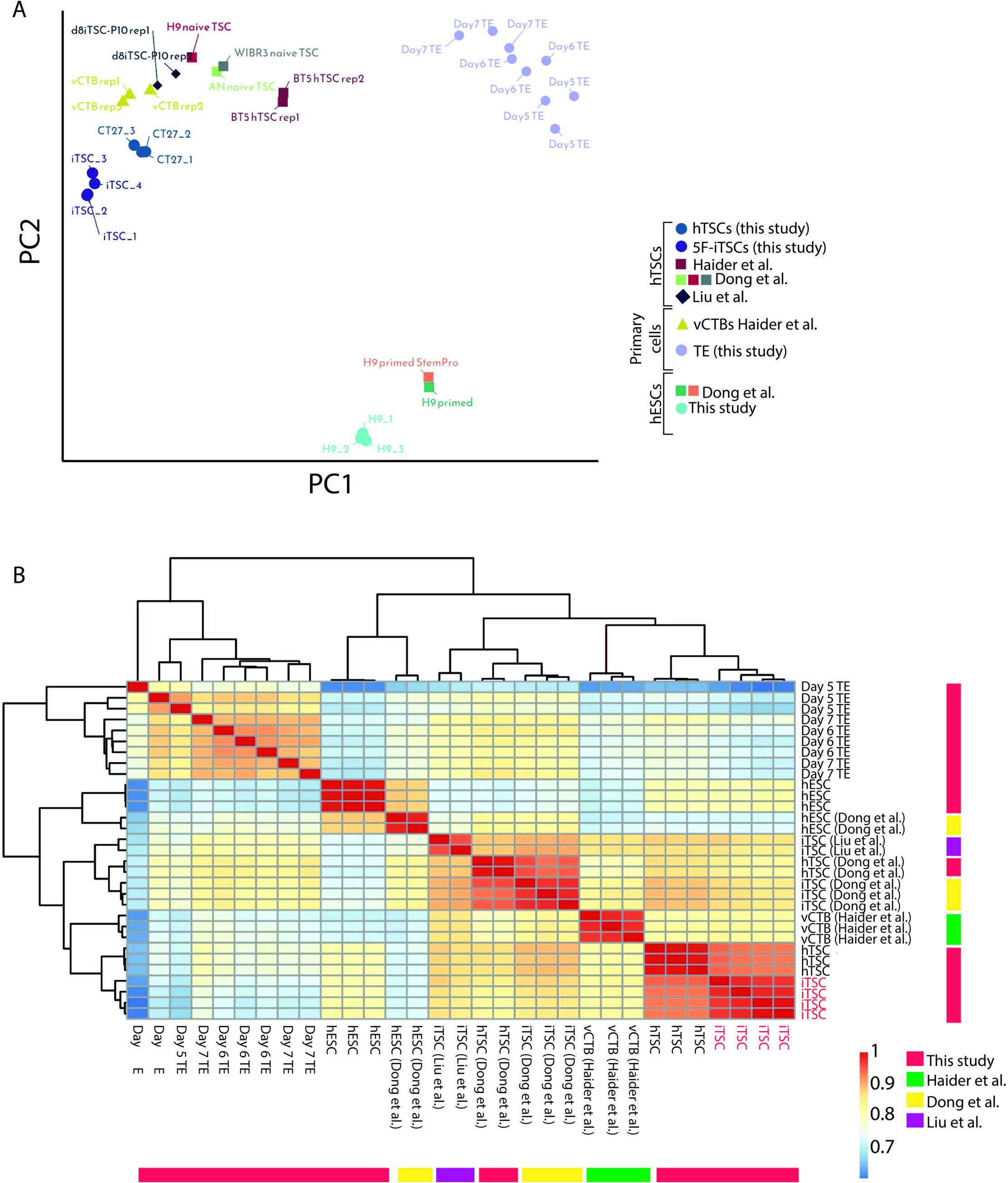
Global transcriptional analysis of iTSCs and similarity to previously established TSCs. Analysis of mRNA-derived iTSCs (this study), previously established hTSCs (CT27, BT5) (Okae et al., 2018), iTSCs derived from naïve transdifferentiation experiments (Dong et al., 2020; Liu et al., 2020), primary placental CTBs (Haider et al., 2018) and primed hESCs (this study)(Dong et al., 2020). (A) Principal component analysis (PCA) using the top 500 most variable expressed genes. (B) Unsupervised hierarchical clustering of the samples. Spearman’s rank correlation coefficient was plotted on a high-to-low scale (red-yellow-blue). iTSCs generated in this study are labelled in red.

### 5F-iTSCs differentiate into both STBs and EVTs

We next examined the differentiation capacity of 5F-iTSCs. We utilised a method to generate terminally-differentiated syncytiotrophoblast as has been previously described (Okae et al., 2018). Accordingly, in both 5F-iTSCs and exisiting hTSCs, we observed formation of multinucleated syncytia which exhibited a reduction in filamentous actin (F-actin) expression, indicating reorganisation of the actin cytoskeleton (Fig. 6A). qRT-PCR of STBs generated from 5F-iTSCs showed significant upregulation of STB-associated genes *ENDOU*, *GCM1*, *PSG3* and *SDC1* compared with undifferentiated 5F-iTSCs (Fig. 6B). hCG is one of the first peptide hormones that is produced by the syncytiotrophoblast (Kliman et al., 1986). Spent culture media from the iTSC-derived syncytiotrophoblast cells was collected after 6 days of differentiation and subjected to an over-the-counter pregnancy test kit which showed detectable hCG expression, similar to exisiting hTSCs (Fig. 6C).

**Figure 6.**
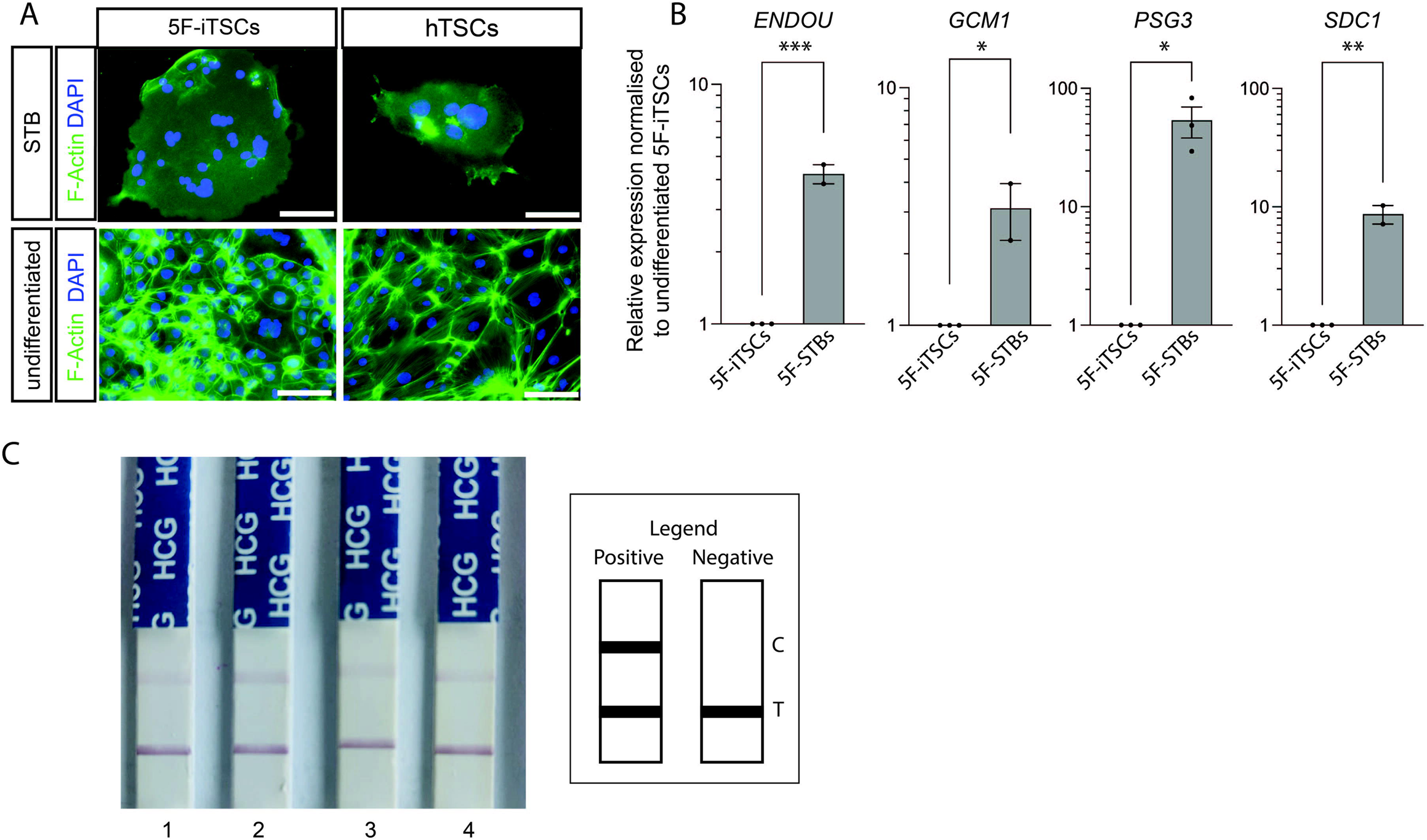
5F-iTSCs can be directed to differentiate to syncytiotrophoblast. (A) Immunostaining analysis of F-actin (green) and DAPI nuclear expression (blue) in STB-differentiated and undifferentiated 5F-iTSCs and hTSCs. Scale bars: 50 µm. (B) qRT-PCR analysis for the detection of STB markers *ENDOU*, *GCM1*, *PSG3* and *SDC1*. Relative expression is shown as fold change over undifferentiated 5F-iTSCs normalised to *GAPDH*. Data are mean ± SEM of n=2 biological replicates analysed with an unpaired one-tailed t-test (*p<0.05, **p<0.01, ***p<0.005). (C) Detection of hCG in spent culture media from syncytiotrophoblast generated from iTSCs and previously established control hTSCs using an over-the-counter pregnancy test. 1. iTSC-derived syncytiotrophoblast replicate 1, 2. iTSC-derived syncytiotrophoblast replicate 2, 3. iTSC-derived syncytiotrophoblast replicate 3, 4. hTSC-derived syncytiotrophoblast.

5F-iTSCs were next differentiated into extravillous trophoblast cells (EVTs) using previously published protocols (Okae et al., 2018). After 6 days we observed morphological changes whereby cells acquired an elongated spindle-like appearance reminiscent of EVTs (Fig. 7A). qRT-PCR analysis of resultant cells revealed significant upregulation of EVT-associated transcripts *HLA-G*, *ITGA1*, *LRRC32*, *LVRN*, *MMP2* and *NOTUM compared to* undifferentiated 5F-iTSCs (Fig. 7B). Altogether this indicates that 5F-iTSCs have the capacity for differentiation to both syncytiotrophoblast and EVT and therefore exhibit all of the hallmarks of hTSCs (Lee et al., 2016; Okae et al., 2018).

**Figure 7.**
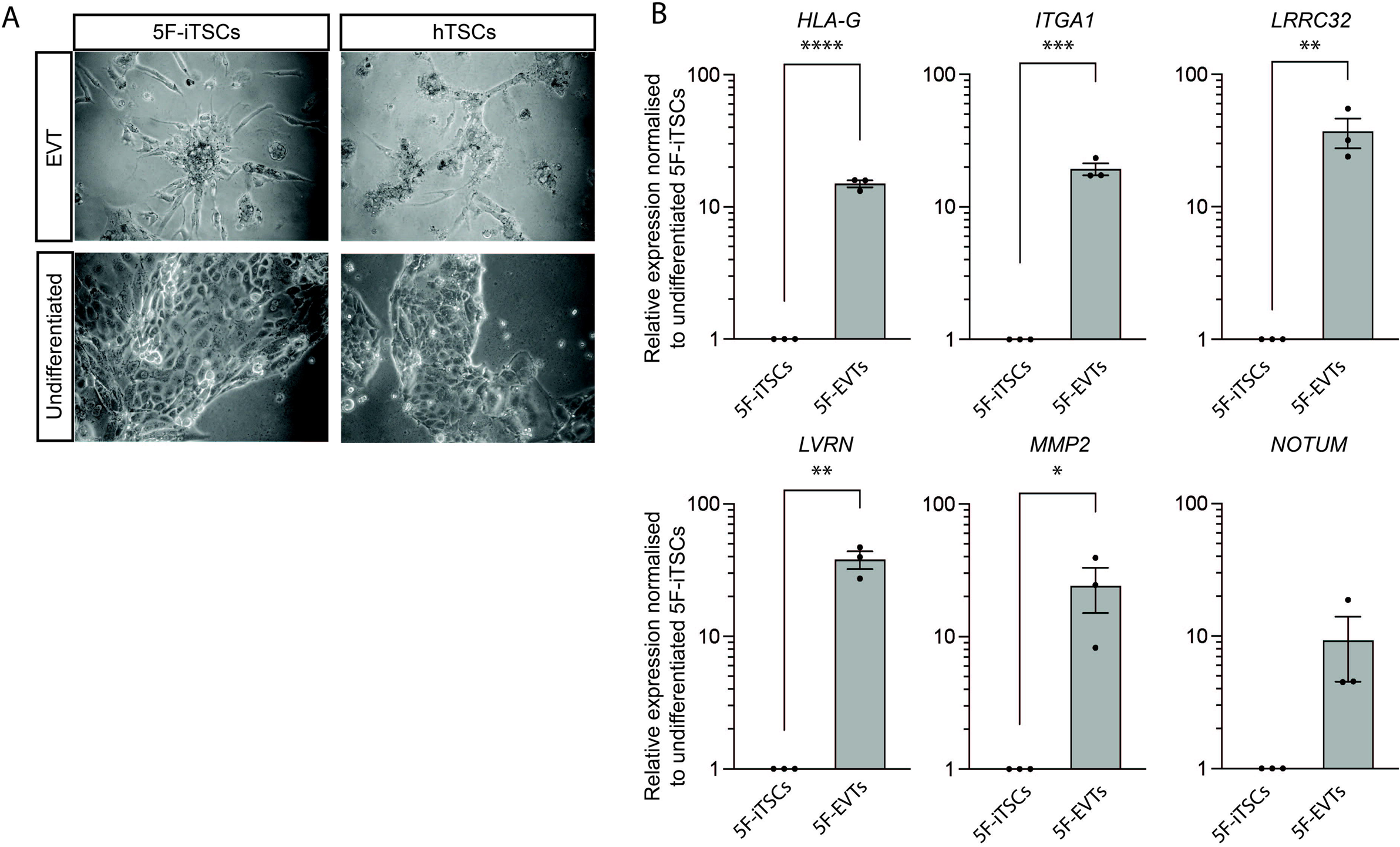
5F-iTSCs can be directed to differentiate to extravillous trophoblast. (A) Brightfield imaging of EVTs differentiated from 5F-iTSCs and hTSCs. (E) qRT-PCR analysis for the detection of selected EVT markers (*HLA-G*, *ITGA1*, *LRRC32*, *LVRN*, *MMP2* and *NOTUM*). Relative expression is shown as fold change over undifferentiated 5F-iTSCs normalised to *GAPDH*. Data are mean ± SEM of n=3 biological replicates analysed with an unpaired one-tailed t-test (*p<0.05, **p<0.01, ***p<0.005, and ****p<0.001).

### Narrowing down transcription factors essential to generate iTSCs

To determine which transcription factors are required for the induction of iTSCs, we examined the effect of withdrawing individual transcription factors from the pool of factors on the formation of 5F-iTSCs. *MYC* was kept constant reasoning that based on its reprogramming effect in other contexts that it functions to generally enhance reprogramming/transdifferentiation rather than being essential for iTSC generation(Nakagawa et al., 2010). To allow us to directly compare the success and efficiency of the iTSC programme induction we maintained an equivalent total amount mRNA in each cocktail by replacing the omitted factors with an equivalent amount of mRNA encoding GFP.

After 10 days of lipofection, cells were immunofluorescently analysed for the expression of two markers of hTSCs, KRT18 and TP63 (Fig. 8A, B; Fig. S2)(Meistermann et al., 2021). Induction of the iTSC-programme was first determined by the presence of cells co-expressing nuclear TP63 and filamentous KRT18 expression extending from the nucleus to the cell membrane, as we observed in cells treated with the 5-factor cocktail (Fig. 8B). Immunofluorescence analysis for the detection of the additional markers, TEAD4 and KRT7, further confirmed the identity of 5F-iTSCs (Fig. S3A). Quantification of immunofluorescence analyses revealed that approximately 80% of cells were KRT18- or TP63-positive when treated with the 5-factor cocktail (Fig. 8C). We observed that only the combination of *GATA2*, *GATA3* and *MYC* produced iTSC colonies, albeit at lower proportions compared to the 5-factor cocktail, with approximately 35% of cells showing co-expression of TP63 and KRT18 (Fig. 8B:10, Fig. 8C). This 3-factor combination led to expression of KRT7 and nuclear TEAD4 (Fig. S3K; yellow arrowheads), similar to those generate in the control 5 factor combination (Fig S3A). By contrast, while the other conditions showed some upregulation of KRT7 and KRT18, the expression pattern of the latter was patchy and disorganised, and the cells did not co-express TP63 nor exhibit nuclear TEAD4 (Fig. 8B; Fig. S3). Next we sought to determine if stable iTSCs could be generated from these 3 factors. We transfected hESCs for 20 days with the *GATA2*, *GATA3* and *MYC* cocktail and observed appearance of hTSC-like colonies, similar to the 5 factor transdifferentiation. These colonies were manually selected and a homogenous 3F-iTSC line was successfully generated (Fig. 8D). qRT-PCR analysis revealed significant upregulation of markers *GATA2*, *GATA3*, *EGFR*, *ENPEP* and *TP63* in 3F-iTSCs compared with the hESC starting population (Fig. 8E). Immunofluorescence confirmed widespread expression of KRT18, TFAP2C and GATA3 (Fig. 8F). However, these 3F-iTSCs could not be maintained long-term in culture, with cell degeneration, loss of colony integrity and loss of proliferation observed after 5 passages. This suggests that *GATA2* and *GATA3* are required for induction of the trophoblast network, but additional factors are required to maintain the cells in long term culture.

**Figure 8.**
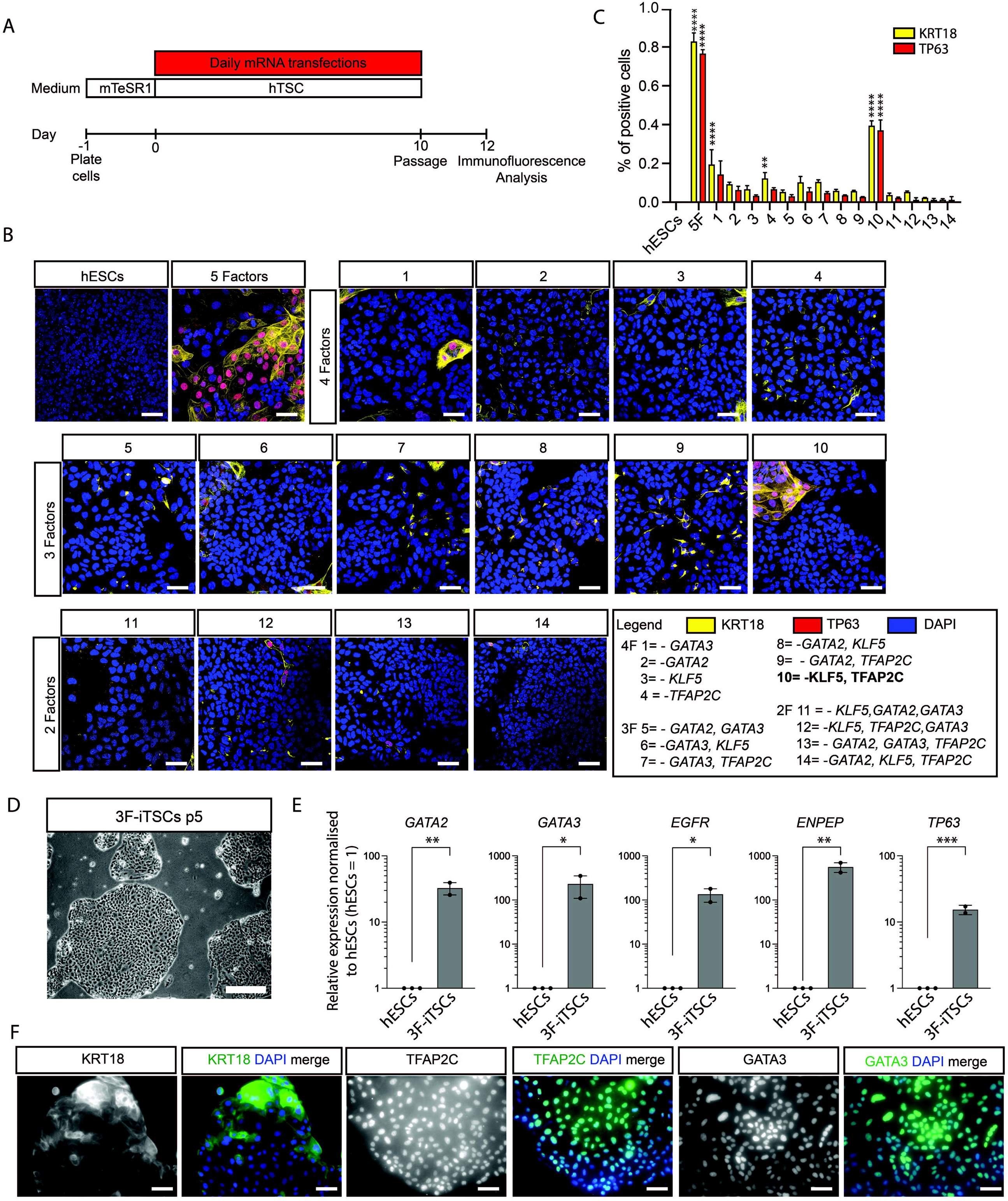
GATA2 and GATA3 are required and sufficient for induction of hTSC programme. (A) Schematic representation of the strategy for testing factors required for iTSC transdifferentiation. hESCs were transfected with cocktails of modified mRNAs where omitted factors were replaced with equivalent amount of mRNA encoding GFP to keep the molar ratio consistent. Cells were transfected daily for 10 days at which point they were fixed for immunofluorescence analysis. (B) Immunofluorescence analysis for the detection of TP63 (red) and KRT18 (yellow) and DAPI nuclear expression (blue) in transfected cells. Merged images are shown. Key for mRNA cocktail combinations: 1, *GATA2, TFAP2C, KLF5* and *MYC* (*GATA3* omitted); 2, *GATA3*, *TFAP2C*, *KLF5* and *MYC* (*GATA2* omitted); 3, *GATA2*, GAT*A3*, *TFAP2C* and *MYC* (*KLF5* omitted); 4*, GATA2*, *GATA3*, *KLF5* and *MYC* (*TFAP2C* omitted); 5, *TFAP2C*, *KLF5* and *MYC* (*GATA2* and *GATA3* omitted); 6, *GATA2*, *TFAP2C* and *MYC* (*GATA3* and *KLF5* omitted); 7, *GATA2*, *KLF5* and *MYC* (*GATA3* and *TFAP2C* omitted); 8, *GATA3*, *TFAP2C* and *MYC* (*GATA2* and *KLF5* omitted); 9, *GATA3*, *KLF5* and *MYC* (*GATA2* and *TFAP2C* omitted); 10, *GATA2*, *GATA3* and *MYC* (*TFAP2C* and *KLF5* omitted); 11, *TFAP2C* and *MYC* (*KLF5*, *GATA2* and *GATA3* omitted); 12, *GATA2* and *MYC* (*GATA3*, *TFAP2C* and *KLF5*); 13*, KLF5* and *MYC* (*GATA2*, *GATA3* and *TFAP2C* omitted) and 14, *GATA3* and *MYC* (*GATA2*, *KLF5* and *TFAP2C* omitted). Scale bars: 50 µm. (C) Quantification of KRT18- and TP63-positive cells. Data is mean ± SEM analysed with analysed with an unpaired t-test (*p<0.05, **p<0.01, and ***p<0.005) (D) Brightfield imaging of 3F-iTSCs line derived and maintained in culture for 5 passages. (E) RT-qPCR analysis for the detection of *GATA2*, *GATA3*, *ENPEP*, *EGFR* and *TP63* in 3F-iTSCs. Relative expression is reflected as fold change over hESCs cultured in mTeSR media normalized to *GAPDH.* Data are mean ± SEM, n=2-3 biological replicates analysed with an unpaired one-tailed t-test (*p<0.05, **p<0.01, and ***p<0.005). (F) Immunofluorescence analysis for the detection of selected trophoblast markers KRT18, TFAP2C, and GATA3 (green) and DAPI nuclear expression (blue) in 3F-iTSCs. Scale bar: 50 µm.

## Discussion

In this study we investigate the sufficiency of TE-associated transcription factors in transdifferentiation of hESCs to iTSCs. In the mouse, CDX2, GATA3, TFAP2C and EOMES were identified as key regulators of the TE lineage (Russ et al., 2000; Strumpf et al., 2005) (Kuckenberg et al., 2010; Ralston et al., 2010). These factors are highly enriched in mouse TE and mTSCs and have been shown to be capable of programming fibroblasts towards mouse iTSCs (Benchetrit et al., 2015; Kubaczka et al., 2015). In this study we sought to identify human-specific TE transcription factors capable of transdifferentiating primed hESCs to iTSCs.

We identified GATA2, GATA3, TFAP2C and KLF5 as factors that are enriched in the human TE. Together with MYC, these transcription factors were capable of transdifferentiating primed hESCs to iTSCs that were transcriptionally equivalent to existing hTSCs. In support of GATA3 as a candidate hTSC transdifferentiation factor is the recent work implicating GATA3 in the initiation of the TE lineage in human embryogenesis (Gerri et al., 2020). GATA2 is exclusively expressed in the TE of mouse, cow and human embryos, and regulates some trophoblast-related genes in mouse and cow (Bai et al., 2011; Gerri et al., 2020; Ma et al., 1997). The mouse homologue of *TFAP2C* (*Tcfap2c*) is required post-implantation with knockout mice dying at embryonic day 7.5 due to proliferation and differentiation defects in the TE compartment (Auman et al., 2002; Werling and Schorle, 2002). In mouse TSCs, TCFAP2C is detected at the promoters of key genes including *Elf5, Gata3, Hand1, Id2* and *Tead4 (Kidder and Palmer, 2010).* By contrast, the role of *TFAP2C* in human trophoblast biology is not fully understood. It is known, however, that TFAP2C is a defining marker of CTBs across gestation (Lee et al., 2016) and it stimulates human placenta lactogen and hCG production (Richardson et al., 2001). Functional analysis of KLF5 in human TE and trophoblast has not yet been performed, but in the mouse this transcription factor is required for the establishment of the ICM and TE. *Klf5*^-/-^ embryos arrest at the blastocyst stage due to defects in TE development resulting in a failure to hatch (Lin et al., 2010). The expression patterns of these factors during human embryogenesis, together with their associated roles in TE and trophoblast biology, implicate *GATA2*, *GATA3*, *TFAP2C* and *KLF5* as candidate regulators of a human TE programme. Indeed whilst GATA2, GATA3 and TFAP2C, along with TFAP2A, have been previously reported to differentiate hESCs to trophectoderm-like cells, these studies were not successful in establishing stable TSC lines (Krendl et al., 2017).

The identity of trophoblast-like cells generated from primed hESCs by modulating the signalling environment has been under investigation for the last two decades (Bernardo et al., 2011; Roberts et al., 2018; Xu et al., 2002). More recently it has been shown that primed hESCs treated with BMP, A83-01 and PD173074 acquire a amnion-like properties, in contrast to naïve cells which in the same conditions generate hTSCs (Io et al., 2021). Another study demonstrated that while a 48 h pretreatment of primed hESCs with BMP4 before culturing in hTSC media generate hTSC-like cells these cells cannot be maintained in culture, nor be directed to differentiate to STB and EVT (Kobayashi et al., 2022). Significantly here we demonstrate the first successful generation of iTSCs from primed hESCs that exhibit transcriptional and epigenetic properties consistant with hTSCs, and can be maintained long-term in culture and have bipotent differentiation capabilities.

Other groups have reported that resetting hESCs from primed to naïve pluripotency generates a small side-population of hTSCs that can be propagated in culture and recapitulate established hTSCs in terms of molecular markers, transcriptome, and directed differentiation potential (Castel et al., 2020; Cinkornpumin et al., 2020; Dong et al., 2020; Liu et al., 2020). Additionally, naïve hESCs treated with either small molecule inhibitors of ERK/mitogen-activated protein kinase (MAPK) and Nodal signalling for 3-5 days (Guo et al., 2021), or for 3 days with transient addition of BMP4 and JAK1 inhibitor (Io et al., 2021), differentiate to hTSCs when placed in conventional hTSC media. This was in contrast to primed hESCs which, in the same conditions, generated amnion-like cells, leading to the hypothesis that primed hESCs had lost the developmental potential to make trophectoderm (Guo et al., 2021). Human naïve ESCs represent an earlier, less-fixed developmental state that reflect gene expression and epigenetic profiles similar to those of the early epiblast or late morula (Gafni et al., 2013; Takashima et al., 2014; Theunissen et al., 2014). A feature of naïve pluripotency is the expression of TFAP2C. TFAP2C plays a critical role during primed to naïve reversion by facilitating the opening of naïve-specific enhancers, as well as regulating the expression of the pluripotency factors OCT4 (Pastor et al., 2018) and KLF4 (Chen et al., 2018). Indeed, CRISPR/Cas9-mediated knockout of *TFAP2C* in naïve hESCs significantly affected the efficiency of transdifferentiation to hTSCs (Guo et al., 2021). This suggests a dual functionality of TFAP2C in human embryogenesis and stem cell counterparts in terms of maintaining naïvety and regulating a TE transcriptional network. Indeed, TFAP2C is suggested to both activate and repress target genes (Eloranta and Hurst, 2002; Pastor et al., 2018). Further molecular analysis of hTSCs revealling the TE-specific repertoire of TFAP2C gene targets may provide further insight into the molecular mechanisms underpinning the TE transcriptional network. Moreover, further detailed characterisation of cells undergoing transdifferentiation to iTSCs would be informative to determine whether primed hESCs transdifferentiate directly to iTSCs or alternatively transit through an amnion-like state or via a reversion to naïve hESCs before committing to a TSC state.

Interesting, all *in vitro* derived TSCs, including the 5F-iTSCs generated in this study more closely resemble the post-implantation CTB cells rather than the preimplantation TE. This suggests that the *in vitro* conditions currently used diverge from the preimplantation *in vivo* niche. Elucidating siganalling pathways active in the epiblast during human embryogenesis has led to the refinement of more physiological culture conditions and established human embryonic stem cell lines more closely resembling their *in vivo* counterparts (Wamaitha et al., 2020). Further investigations of human TE and systematic evaluation of culture conditions may similarly lead to the derivation of TSC lines that more closely resemble this earlier TE developmental timepoint.

The combinatorial transdifferentiation experiments provided further insights in the roles of TE-associated transcription factors. Our 3-factor experiments suggest that both *GATA2* and *GATA3* are essential for upregulation of the hTSC programme. The GATA family of transcription factors are implicated in establishing cell fate during development and often show functional redundancy between members (Fujiwara et al., 2004; Peterkin et al., 2005). However it is unclear whether GATA2 and GATA3 play different roles here in transdifferentiation, or if there is functional redundancy between the two. Interestingly in our time-course analysis of transdifferentiation, the timing of onset of endogenous *GATA2*, as well as the levels to which it is upregulated, appear to lag that of *GATA3* expression. This may indicate that while both are required, GATA3 acts upstream of GATA2. Future experiments including epistatic functional studies and investigation of transcription factor binding and chromatin occupancy to distinguish the roles of these factors would elucidate these mechanisms. Surprisingly, and in contrast to the mouse, *TFAP2C* appears dispensable for transdifferentiation. While our transdifferentiation strategies were successful in establishing stable lines, in the future the efficiency of hTSC transdifferentiation may be further enhanced by either equivalently increasing the amount of each factor or by altering the stoichiometry of transcription factors, as has been demonstrated for induced pluripotent stem cell and cardiomyocyte reprogramming (Carey et al., 2011; Muraoka and Ieda, 2015). The mRNA-based strategy is advantageous to address this as it more easily allows for the generation of bespoke combinations of transcription factor as well as the alteration of the stoichiometry of factors in the cocktail, compared to viral based methods. In addition, functional examination of these factors in knock-out hTSCs may further refine the combination of essential factors required for the maintenance of these cell types.

Defective TE specification, trophoblast differentiation and maturation leading to abnormal placentation underlies miscarriage and preeclampsia (Burton and Jauniaux, 2017; Fisher, 2015). hTSCs and trophoblast organoids present a paradigm for modelling placenta development and disease *in vitro*. However, a caveat of existing CTB- and blastocyst-derived hTSCs and trophoblast organoids is that it cannot be ascertained whether the starting population of cells would have given rise to a normal or a disease-affected placenta. Initial attempts to derive hTSCs and trophoblast organoids directly from placental tissue after 12 weeks’ gestation, or at even later stages coincident with placental-related disease manifestation, were unsuccessful (Haider et al., 2018; Okae et al., 2018; Turco et al., 2018). More recently it has been suggested that term CTB cells could be maintained in conventional TSC media under hypoxic conditions through the repression of regulator of trophoblast differentiation GCM1 (Wang et al., 2022). In addition, iTS-like cells can be generated from term CTB cells by overexpression of *TFAP2C*, *TEAD4*, *CDX2*, *ELF5*, and *ETS2* (Bai et al., 2021). However in both these studies the cells were not assessed for hallmarks of trophoblast including *ELF5* promoter methylation status and C19MC miRNA expression and the cells remain distinct from the primary term CTB cells from which they were derived.

We lack an understanding of the genetic basis of placental-related diseases and so we cannot currently apply CRISPR/Cas9-mediated genome editing to create mutations for disease modelling in existing hTSCs. Instead, we propose that the mRNA transdifferentiation strategy presented here could be applied and further refined to generate iTSCs from patient fibroblasts, or fibroblasts or mesenchymal stromal cells isolated from disease-affected placentas (Pelekanos et al., 2016). This strategy could open up the possibility of generating patient-specific human TSCs. Indeed, we have attempted to apply the strategy described here to fibroblasts but our pilot studies were unsuccessful. This may suggest additional transcription factors may be required for fibroblasts to make the conversion. Future experiments may include overexpression of NR2F2 as our time-course analysis suggests that it is one of the most highly upregulated transcription factors during transdifferentiation. Alternatively, if fibroblasts are not amenable to this conversion, reprogramming could be first employed to generate patient-specific primed iPSCs as the starting population before applying our five-factor strategy. This would have an advantage over transiting through naïve iPSCs if it is found that epigenetic imprints are lost during iTSC generation depending on the starting cell type or if there are persistent issues of karyotypic instability in the starting cell type. Additionally, mRNA transfection has the benefit of being readily transferable to future clinical application as it avoids the random integration of transgenes that can result in genomic modification and tumorigenicity (Warren et al., 2010). In all, this strategy could allow for the generation of a catalogue of clinically normal and disease-associated hTSCs that would provide a tool for basic research into trophoblast biology as well as a powerful tool for elucidating placental defects including recurrent miscarriage, preeclampsia, intrauterine growth restriction and stillbirth as well as a future drug screening platform.

## Methods

### Human embryo thaw and culture conditions

Human embryos that were surplus to family building requirements were donated to the Francis Crick Institute for use in research projects under the UK Human Fertilisation and Embryology Authority License number R0162 and the Health Research Authority’s Research Ethics Committee (Cambridge Central reference number 16/EE/0067). Slow-frozen blastocysts (day 5 and day 6) were thawed using the BlastThaw (Origio; Cat. No. 10542010A) kit using the manufacturer’s instructions. Vitrified blastocysts (day 5 and day 6) were thawed using the vitrification thaw kit (Irvine Scientific; Cat. No. 90137-SO) following the manufacturer’s instructions. Human embryos were cultured in pre-equilibrated Global Media supplemented with 5 mg/ml Life Global HSA (both LifeGlobal; Cat. No. LGG-020 and LGPS-605) and overlaid with mineral oil (Origio; Cat. No. ART-4008-5P) and incubated in Embryoscope+ time lapse incubator (Vitrolife). Embryos were grouped into day 5, 6 or 7 based on data provided from the clinc. For the collection of day 5 samples, embryos were fixed approximately 2 h after thawing to allow them to recover. To collect day 6 and 7 samples, day 5 or 6 embryos were cultured for the appropriate time before fixation.

### Microdissection of TE from human blastocysts

Embryos were placed in drops of G-MOPS solution (Vitrolife; Cat. No. 10129) on a petri dish overlaid with mineral oil. The plate was placed on a microscope stage (Olympus IX70) and the embryos were held with an opposing holding pipette and blastomere biopsy pipette (Research Instruments) using micromanipulators (Narishige, Japan). The biopsy mode of a Saturn 5 laser (Research Instruments) was used to separate the mural TE from the ICM and polar TE. The separated mural TE was transferred to individual low bind RNAse-free tube containing 0.25 μl RNase inhibitor, 4.75 μl dilution buffer (SMARTer Ultra Low Input RNA kit, Clontech; Cat. No. 634820) and 5 μl nuclease-free water on a pre-chilled CoolRack (Biocision, CA). Samples were stored at −80°C until ready to be processed.

### cDNA synthesis and library preparation of TE samples

cDNA was synthesized using SMARTer Ultra Low Input RNA for Illumina Sequencing-HV kit (Clontech Laboratories; Cat. No. 634820) according to the manufacturer’s instructions and as previously published (Blakeley et al., 2015; Hyslop et al., 2016). cDNA was sheared using Covaris S2 with the modified settings 10% duty, intensity 5, burst cycle 200 for 21min. Libraries were prepared using Low Input Library Prep Kit (Clontech Laboratories; Cat. No. 634900) according to the manufacturer’s instructions. Library quality was assessed with an Agilent 2100 BioAnalyser and concentration measured by QuBit broad range assay (Thermo Fisher; Cat. No. Q32850). Prepared libraries were submitted for 50-bp paired-end sequencing on Illumina HiSeq 2000.

### cDNA synthesis and library preparation of bulk cell lines

For bulk RNA-seq of cell lines, RNA was isolated using TRI reagent (Sigma-Aldrich; Cat. No. T9424) and DNase I-treated (Ambion; Cat. No. AM2222). Libraries were prepared using the polyA KAPA mRNA HyperPrep Kit (Roche; Cat. No. 8098115702). Quality of submitted RNA samples and the resulting cDNA libraries was determined by ScreenTape Assay on a 4200 TapeStation (Agilent). Prepared libraries were submitted for single-end 75 bp sequencing on an Illumina HiSeq 4000 (Illumina).

### RNA-seq analysis of TE samples

RNA-seq data from human TE was analysed as previously described (Blakeley et al., 2015; Hyslop et al., 2016). Briefly, the reference human genome sequence was obtained from Ensembl, along with the gene annotation (GTF) file. The reference sequence was indexed using the bowtie2-build command. Reads were aligned to the reference human genome sequence using Tophat2 (Kim et al., 2019), with gene annotations to obtain BAM files for each sample. BAM files were then sorted by read coordinates and converted into SAM files using SAMtools. The process of mapping and processing BAM files was automated using a custom Perl script. The number of reads mapping to each gene were counted using the program HTSeq-count (Version 0.6.1; (Anders et al., 2015). The resulting count files for each sample were used as input for differential expression analysis using DESeq2 (Anders and Huber, 2010). Firstly, the function ‘estimateSizeFactors’ and ‘estimateDispersions’ were used to estimate biological variability and calculate normalised relative expression values across the different blastocyst samples. Initially, this was performed without sample labels (option: method=’blind’) to allow unsupervised clustering of the blastocyst samples using principal components analysis and hierarchical clustering. The dispersion estimates were recalculated with the sample labels included and with the option: method=’pooled’. The function ‘nbinomTest’ was then used to calculate p-values to identify genes that show significant differences in expression between different developmental stages.

An RPKM >5 threshold was applied to generate the lists of genes expressed at each stage analysed. Overlap between the gene lists was determined using the online web application GeneVenn(Pirooznia et al., 2007). Gene lists were used to perform a Gene Ontology and REACTOME functional enrichment analysis (Ashburner et al., 2000; Fabregat et al., 2018) to identify overrepresented categories using a significance threshold of p ≤0.05.

### RNA-seq data analysis of induced trophoblast stem cells (iTSCs) and related lines Data from this study

Sequencing data from sample replicates comprising iTSCs (n = 4), H1 ES cells (n = 3) and primary CT27 cells (n = 3) were first checked using the *FastQC* package (https://www.bioinformatics.babraham.ac.uk/projects/fastqc/). Adapter removal was performed using *Trimgalore v0.6.6* (https://github.com/FelixKrueger/TrimGalore), and the trimmed read data were re-checked for conformity and quality with *FastQC*.

### Accessory datasets

Primary cytotrophoblast gene expression data was downloaded from GEO with accession number GSE109976 (Haider et al., 2018). Additional primed hESCs gene expression data was downloaded from GEO with accession numbers GSM4116153 and GSM4116151 (Dong et al., 2020). Trophoblast stem cells derived from human blastocysts gene expression data was downloaded from GSE138762 (Dong et al., 2020). Trophoblast stem cells derived from fibroblast reprogramming gene expression data was downloaded from GSE150616 (Liu et al., 2020) and GSE138762 (Dong et al., 2020)

Next, sequences were aligned to the reference genome (Homo_sapiens.GRCh38) using *HISAT2 v2.2.1* (Kim et al., 2019), with trimming of 5’ and 3’ bases performed based on the QC of each of each of these samples. Counts were generated using *FeatureCounts v2.0.1*, and the count matrix was analysed further using *DESeq2* (Liao et al., 2019; Love et al., 2014). Genes with fewer than 10 counts detected across all 22 samples were excluded, leaving 17498 genes for further processing. A variance stabilizing transformation (VST) was applied, and subsequently batch related effects in combining these different datasets were corrected for using the *limma* package (i.e. *limma::removeBatchEffect)*(Law et al., 2016). Sample to sample distances were computed and plotted as a heatmap. Principal Component Analysis (PCA) was performed using the top 500 most highly variable genes.

### Generation of inducible system

Doxycycline-inducible overexpression of transcription factors was achieved using the Lenti-X Tet-On 3G Inducible Expression System (Clontech; Cat. No. 631363) following the manufacturer’s protocol, and as previously described (Wamaitha et al., 2015). Coding sequences were sub-cloned from the template plasmids into the pLVX-TRE3G vector to generate individual pLVX-TRE3G-Gene of Interest (GOI) vectors (plasmids available upon request). Lentiviral packaging was achieved using 7 μg of pLVX-TRE3G-GOI and the Lenti-X Packaging Single Shot reagents in HEK-293T cells. Transfection media was replaced after 6 h with fresh MEF media. Lentiviral supernatants were subsequently collected after transfection of HEK 293T cells with either the pLVX-TRE3G-GOI or the pLVX-Tet3G vector and concentrated using X-fect reagent (Clontech; Cat. No. 631317). Supernatants were concentrated by ultracentrifugation. Equal volumes of lentivirus supernatant encoding *GATA2*, *GATA3*, *TFAP2C*, *MYC* and *KLF5* were pooled to generate a 5-factor cocktail which was aliquoted into single-use 10 µl volumes. hESCs were grown to 70% confluency and transduced with the pLVX-tet3G lentivirus followed by selection with G418 (250 μg/ml) for one week. G418-resistant cells were selected for at least 2 passages and then transduced with the pLVX-TRE3G-GOI lentivirus pool, followed by selection with puromycin (0.5 μg/ml) for four days. For the induction of GOI expression, doxycycline was added to the mTeSR1 media at a concentration of 1 μg/ml. Clonal lines were generated and screened for transgene integration. Transgene integration was confirmed by culturing clonal lines in the presence of doxycycline in mTeSR for 48 h and transgene expression was detected by RT-qPCR using primers distinguishing exogenous expression from endogenous expression, and immunofluorescence analysis for TFAP2C due to difficulties in designing primers to distinguish between exogenous and endogenous transcripts. Primer sequences used to detect transgene expression are detailed in Table S5.

### Generation of templates for lentivirus production and in vitro transcription (IVT)

The vector design strategy was informed from previously published reports (Mandal and Rossi, 2013). Briefly, nucleotide sequences of the open reading frames for the canonical isoforms of transdifferentiation factors were identified from the Ensembl genome browser, and the sequences were verified against the corresponding amino acid sequences in Uniprot. Individual *in vitro* transcription (IVT) template constructs for *G A T A 2, G A T A 3, TFAP2C* and *K L F 5*consisting of a T7 promoter-51UTR-Open Reading Frame-31UTR-T7 terminator cassette cloned into a pUC57 backbone were custom made (Genewiz UK Ltd). 5’UTR and 3’UTRs were added to maximize stability of mRNA transcripts and to increase protein translation. The ORFs of *M Y C*and *G F P*were templated from plasmids bearing human MYC and GFP and ligated into the pUC57 backbone flanked by the 5’UTR and 3’UTR sequences. Annotated sequence files of all constructs are provided in Table S7.

### Generation of modified mRNAs by in vitro synthesis

The mRNA synthesis protocol has been described previously (Mandal and Rossi, 2013). Briefly, dsDNA templates were linearised from cDNA clones in pLVX vectors for *GATA2, GATA3, TFAP2C, KLF5* and *MYC*. A small amount of digestion mix was run on a gel to confirm complete digestion. Linearised plasmid was purified using PCR purification kit (Qiagen; Cat. No. 28104). A nanodrop was used to confirm the purity of the eluted product according to the 260/280 ratios. Poly(A) tail was added using KAPA PCR ready mix (2X), Xu-F1 and Xu-T120 primers (Integrate DNA Technologies) and digested plasmid adjusted to 10 ng/ul. Tail PCR was run for 32 cycles and purified using PCR purification kit. *In vitro* transcription was performed using MEGAscript T7 kit (Thermo Fisher; Cat. No. AMB13345): custom NTP mix was prepared with 3’-O-Me-m7G cap analogue (60 mM, NEB), GTP (75mM, MEGAscript T7 kit), ATP (75mM, MEGAscript T7 kit), Me-CTP (100 mM, TriLink; Cat. No. N-1014-1) and pseudo-UTP (100mM, Tri-link; Cat. No. O-0263). Reaction was heated at 37 C for 2 h. 2 ul of Turbo DNase (Thermo Fisher; Cat. No. AM2238) was added and incubated at 37 C for 15 min. DNAse treated reaction mix was purified using RNAeasy kit (Qiagen; Cat. No. 74104) according to the manufacturer’s instructions. RNA was phosphatase-treated using Antarctic phosphatase (New England BioLabs; Cat. No. M0289S) and purified using MEGAclear kit (Thermo Fisher; Cat. No. AM1909). Concentration and quality were measured using a nanodrop and adjusted to 100 ng/ul. For transdifferentiation experiments a modified mRNA cocktail was prepared by mixing the five factors in an equal molar ratio. The cocktail was aliquoted into single use aliquots containing a total mRNA amount of 1 µg and stored at −80 C until needed. For transcription factor combinatorial experiments cocktails were made up to a total amount of 500 ng of mRNA with the relevant quantity of mRNA encoding GFP replacing the omitted factors.

### Cell culture

H1 hESCs (WiCell) cells were routinely cultured in mTeSR1 (Stem Cell Technologies; Cat. No. 85850) and growth factor reduced Matrigel (BD Biosciences; Cat. No. 356321). For hTSC transdifferentiation experiments, H1 cells were plated in mTeSR1 media on Matrigel-coated cell culture dishes. Media was changed for hTSC media the next day. hTSC media was prepared as described previously (Okae et al., 2018) (Chir99021, BioTechne, Cat. No. 4423/10; EGF, Tebu Bio Ltd, Cat. No. 167AF-100-15-a; ITS-X supplement, Fisher Sci, Cat. No. 10524233; L-ascorbic acid, Tocris, Cat. No. 4055/50; A83-01, Sigma-Aldrich Cat. No. SML0788/5MG; SB431542, Cambridge Bioscience, Cat. No. SM33-2; Valproic acid, Sigma-Aldrich Cat. No. V-006-1ML; Y27632, Stem Cell Technologies, Cat. No. 72302). For transgene induction, doxycycline was added daily at a concentration of 1 μg/ml over 20 days for transgene induction. hTSCs were passaged weekly by dissociation with TrypLE Express (Fisher Thermo Scientific, Cat. No. 12604013) for 15 min at 37 C and passaged en masse initially onto collagen IV-coated (5 µg/ml; Corning, Cat. No. 354233) 10 cm plates and subsequently onto 6-well plates and maintained in hTSC media. CT27 and BTS5 hTSCs (Riken BRC) were routinely cultured in hTSC media described above on collagen IV-coated plates. Cells were routinely tested for mycoplasma contamination using PCR-based detection assays.

### Lipofection of hESCs

For one well of a 6-well plate, 1 µg (1x) of mRNA was mixed with 86 µl of OptiMEM reduced serum media in an Eppendorf tube. In a second tube 0.5 x volume of lipofectamine RNAi max (Thermofisher; Cat. No. 13778100) was mixed with 93 µl OptiMEM. Tubes were incubated at room temperature for 5 min. The tubes were mixed together and incubated at room temperature for 20 min. The lipofection mix was added to the well in a dropwise manner and mixed well by rocking the plate from side to side. The cells were incubated at 37 C in 20% O_2_ for 4 h. The media was then replaced with hTSC media containing 200 ng/ml recombinant B18R to prevent gamma-interferon response (Stem Cell Technologies; Cat. No. 78075). For transdifferentiation experiments lipofection was performed at the same time every day for 20 days. Mock transfection was performed using the an equivalent amount of *GFP* mRNA for 20 days. For transcription factor combinatorial experiments hESCs were lipofected daily for 10 days with a mRNA transfection cocktail totalling 500ng, were individual factors were replaced with an equivalent amount of *GFP* mRNA.

### Directed differentiation

For the induction of syncytiotrophoblast, 1 x 10^5^ 5F-iTSCs or hTSCs were seeded in hTSC media in a 12-well plate pre-coated with 2.5 mg/ml Col IV. Cells were allowed to adhere for a minimum of 6 h before media was changed to STB media comprised of DMEM/F12 supplemented with 0.1 mM 2-mercaptoethanol, 0.5% penicillin-streptomycin, 0.3% BSA, 1% ITS-X supplement, 2.5 mM Y27632, 2 mM forskolin, and 4% KSR]. The media was replaced at day 3, and resultant cells were analysed at day 6.

For induction of extravillous trophoblast cells, 0.75 x 10^5^ 5F-iTSCs and hTSCs were seeded in hTSC media in 12-well plates pre-coated with 1 mg/ml Col IV. Cells were allowed to adhere for a minimum of 6 h before media was changed to EVT1 media comprised of DMEM/F12 with 0.1 mM 2-mercaptoethanol, 0.5% Penicillin-Streptomycin, 0.3% BSA, 1% ITS-X supplement, 100 ng/ml NRG1, 7.5 μM A83-01, 2.5 μM Y27632, and 4% KSR. Cold growth factor reduced matrigel was added to yield a final concentration of 2%. After 72 h, media was replaced with EVT2 media that lacks NRG1, and matrigel was added to yield a final concentration of 0.5%.

### Immunostaining

Embryos were fixed with 4% paraformaldehyde in PBS for 1 h at 4°C. Immunofluorescence staining was performed as described previously (Fogarty et al. 2018). The primary antibodies used are listed in Table S5. Embryos were placed on µ-Slide 8 well coverslip dishes for confocal imaging (Ibidi; Cat. No. 80826). Imaging was performed using a Leica SP5 confocal microscope and 3 μm thick optical sections were collected. hESC, iTSC and hTSC lines were fixed with PBS 4% PFA for 1 h at 4°C, then washed three times with PBS (Life Technologies; Cat. No. 14190-094). Blocking was achieved by incubation with PBS with 10% donkey serum (Sigma-Aldrich; Cat. No. D9663) and 0.1% Triton X-100 (Sigma-Aldrich; Cat. No. T8787) for 30 mins at room temperature. Permeabilisation was performed by incubation in PBS with 0.5% Triton X-100. Primary antibodies were diluted in blocking solution and each incubated overnight at 4 C. Following each incubation, the cells underwent 3 x 5 min washes with PBS with 0.1% Triton X-100. Secondary antibodies were diluted 1:300 in PBS with 0.1% Triton X-100 for 1 h at room temperature. After 3 x 5 min washes in PBS with 0.1% Triton X-100, 5 μg/ml DAPI (Sigma-Aldrich; D9542) was added for 2 mins during the final wash in order to perform nuclear staining. Confocal imaging analysis of embryos was performed using a Leica SP5. Fluorescent imaging of cells was performed using an Olympus IX73 using CellSens software (Olympus) and a Zeiss Meta LSM510 confocal and Zen Pro 2102 software. For quantification of immunostaining analysis of 2D cells, an average of 250 (Figure 8) or 150 (Figure S3) cells from 5 random frames of view were counted from two independent experiments.

### Quantitative RT-PCR

RNA was isolated using TRI Reagent (Sigma) and DNaseI treated (Ambion). cDNA was synthesised using a Maxima First Strand cDNA Synthesis Kit (Fermentas). Quantitative RT-PCR (qRT-PCR) was performed using Sensimix SYBR Low-Rox kit (Bioline, QT625) on a BioRad CFX96 Touch Real-Time PCR Detection System. Primer pairs (listed in Table S6) were either previously published, as referenced, or designed using Primer3 software. qRT-PCR for the detection of miRNAs was performed using primers and protocols as previously published (Lee et al., 2016). cDNA was synthesised using the TaqMan™ MicroRNA Reverse Transcription Kit (Applied Biosystems, 4366597). qRT-PCR was performed on a CFX384 Touch Real-Time PCR Detection System (Bio-Rad, 1855484) and CFX Maestro software. Statistical analysis was performed using Prism 10 (Graphpad).

### ELF5 promoter methylation analysis

Bisulfite sequencing of the *ELF5* promoter was carried out as described previously (Bernardo et al, 2011). Briefly, bisulphite conversion was carried out on 400 ng genomic DNA using the EpiTect Bisulfite Kit (QIAGEN) following the manufacturer’s instructions. Nested PCR was carried out on 10% of the eluted DNA to analyse −432/-3bp relative to the *ELF5* transcriptional start site. Purified PCR products were cloned using the pGEM-T Easy Vector System (Promega) and sequenced.

## Acknowledgements

We are very grateful to the donors of human embryos whose contributions enable this research. We thank all members of the Niakan and Fogarty labs for help and comments on the paper. We are grateful to the Francis Crick Institute’s Science Technology Platforms: Lyn Healey from Human Embryonic Stem Cell Facility, Robert Goldstone and Deb Jackson from the Advanced Sequencing Facility; and the Advanced Light Microscopy Facility.

## Funding

Work in the laboratory of KKN was supported by the Wellcome 221856/Z/20/Z and by the Wellcome Human Developmental Biology Initiative 215116/Z/18/Z. Work in the K.K.N. laboratory was also supported by the Francis Crick Institute, which receives its core funding from Cancer Research UK (FC001120), the UK Medical Research Council (FC001120), and the Wellcome Trust (FC001120). The M.S. lab at Centre for Gene Therapy and Regenerative Medicine, King’s College London was supported by a Wellcome Trust Clinical Career Development Fellowship (222052/Z/20/Z), and the M.S. group in Singapore (including P.M.) was supported by a National Medical Research Council, Singapore Clinician Scientist Investigator Award (CSAINV17may001). N.M.E.F was supported by a King’s Prize Fellowship and a UKRI Future Leaders Fellowship. For the purpose of Open Access, the authors have applied a CC-BY public copyright licence to any Author Accepted Manuscript version arising from this submission.

## Competing Interests

No competing interests declared.

## Author Contributions

N.M.E.F and K.K.N. conceived the study; K.K.N and N.M.E.F supervised the project; N.M.E.F and K.K.N designed the experiments. N.M.E.F., P.A.B., and A.A. performed the majority of experiments with assistance from A.C., L.D., and A.M. C.E.S. performed *ELF5* methylation analysis. K.K.N. performed embryo dissections. K.E., P.S., L.C. and R.A.O. coordinated the donation of embryos to the research project. P.B., P.M. and M.S. performed bioinformatic analysis of RNA-seq data. N.M.E.F. and K.K.N. wrote the manuscript with help from all the authors.

## Data availability

RNA-seq FastQ files have been deposited at ArrayExpress with the accession numbers E-MTAB-10749 for data from microdissected human embryos and E-MTAB-10748 for data arising from cell lines.

## Notes

### Competing Interest Statement

The authors have declared no competing interest.

### Summary of Updates

We have performed significant additional experiments as follows: 1. We have performed a time-course RT-qPCR analysis during the five factor transdifferentiation. We assessed expression of trophoblast-associated transcripts in induced cells relative to cells grown in hTSC media alone. Here we demonstrate that while at day 15 we get significant upregulation of endogenous GATA3, TP63 and ENPEP, it is at day 20 that we also have significant upregulation of NR2F2 and trophoblast-associated cell surface markers (Figure 3C). This confirms to us that 20 days of overexpression is required to induce the downstream trophoblast network. 2. We confirm that 5F-iTSCs express trophoblast-specific miRNAs from the C19MC cluster, comparable to conventional hTSCs (Figure 4G). 3. We confirm that 5F-iTSCs are hypomethylated at the ELF5 promoter, comparable to conventional hTSCs (Figure 4H). 4. We re-analysed the RNA-seq data and included samples of human trophectoderm. We have re-analysed the data by including the TE samples. We now see that 3 clusters are formed: 1. 5F-iTSCs, primary cytotrophoblast cells, all existing hTSC, 2. Primary TE and 3. hESCs. Accordingly we no longer see this separation on the basis of in vitro/in vivo differences (Figure 5A and B). 5. We performed additional molecular characterisation of 5F-iTSC derived syncytiotrophoblast. We performed qRT-PCR analysis for a panel of STB markers and observed significant upregulation compared to undifferentiation 5F-iTSCs (Figure 6B). 6. We confirm that 5F-iTSCs differentiate to EVTs equivalently to existing hTSCs based on morphology. We also performed qRT-PCR analysis for the detection of markers of EVTs and show significant upregulation compared to undifferentiated 5F-iTSCs (Figure 7). 7. We demonstrate the establishment of stable lines from 3 factor transdifferentiation. While we see characteristic expression of hTSC-markers at the RNA and protein level, these cells cannnot be maintained long term in vitro. We observed cell vaculoation, loss of colony integrity, loss of proliferation, and cells eventually lifted off. Thus we could not test their differentiation potential (Figure 8,D, E and F).

## References

Anders, S., Huber, W. (2010). Differential expression analysis for sequence count data. Genome Biol 11, R106.

Anders, S., Pyl, P.T., Huber, W. (2015). HTSeq--a Python framework to work with high-throughput sequencing data. Bioinformatics 31, 166–169.

Apps, R., Gardner, L., Hiby, S.E., Sharkey, A.M., Moffett, A. (2008). Conformation of human leucocyte antigen-C molecules at the surface of human trophoblast cells. Immunology 124, 322–328.

Ashburner, M., Ball, C.A., Blake, J.A., Botstein, D., Butler, H., Cherry, J.M., Davis, A.P., Dolinski, K., Dwight, S.S., Eppig, J.T., Harris, M.A., Hill, D.P., Issel-Tarver, L., Kasarskis, A., Lewis, S., Matese, J.C., Richardson, J.E., Ringwald, M., Rubin, G.M., Sherlock, G. (2000). Gene ontology: tool for the unification of biology. The Gene Ontology Consortium. Nat Genet 25, 25–29.

Auman, H.J., Nottoli, T., Lakiza, O., Winger, Q., Donaldson, S., Williams, T. (2002). Transcription factor AP-2gamma is essential in the extra-embryonic lineages for early postimplantation development. Development 129, 2733–2747.

Baczyk, D., Dunk, C., Huppertz, B., Maxwell, C., Reister, F., Giannoulias, D., Kingdom, J.C. (2006). Bi-potential behaviour of cytotrophoblasts in first trimester chorionic villi. Placenta 27, 367–374.

Bai, H., Sakurai, T., Someya, Y., Konno, T., Ideta, A., Aoyagi, Y., Imakawa, K. (2011). Regulation of trophoblast-specific factors by GATA2 and GATA3 in bovine trophoblast CT-1 cells. J Reprod Dev 57, 518–525.

Bai, T., Peng, C.Y., Aneas, I., Sakabe, N., Requena, D.F., Billstrand, C., Nobrega, M., Ober, C., Parast, M., Kessler, J.A. (2021). Establishment of human induced trophoblast stem-like cells from term villous cytotrophoblasts. Stem Cell Res 56, 102507.

Bansal, A.S., Bora, S.A., Saso, S., Smith, J.R., Johnson, M.R., Thum, M.Y. (2012). Mechanism of human chorionic gonadotrophin-mediated immunomodulation in pregnancy. Expert Rev Clin Immunol 8, 747–753.

Benchetrit, H., Herman, S., van Wietmarschen, N., Wu, T., Makedonski, K., Maoz, N., Yom Tov, N., Stave, D., Lasry, R., Zayat, V., Xiao, A., Lansdorp, P.M., Sebban, S., Buganim, Y. (2015). Extensive Nuclear Reprogramming Underlies Lineage Conversion into Functional Trophoblast Stem-like Cells. Cell Stem Cell 17, 543–556.

Bernardo, A.S., Faial, T., Gardner, L., Niakan, K.K., Ortmann, D., Senner, C.E., Callery, E.M., Trotter, M.W., Hemberger, M., Smith, J.C., Bardwell, L., Moffett, A., Pedersen, R.A. (2011). BRACHYURY and CDX2 mediate BMP-induced differentiation of human and mouse pluripotent stem cells into embryonic and extraembryonic lineages. Cell Stem Cell 9, 144–155.

Blakeley, P., Fogarty, N.M., del Valle, I., Wamaitha, S.E., Hu, T.X., Elder, K., Snell, P., Christie, L., Robson, P., Niakan, K.K. (2015). Defining the three cell lineages of the human blastocyst by single-cell RNA-seq. Development 142, 3151–3165.

Burton, G.J., Jauniaux, E. (2017). The cytotrophoblastic shell and complications of pregnancy. Placenta 60, 134–139.

Carey, B.W., Markoulaki, S., Hanna, J.H., Faddah, D.A., Buganim, Y., Kim, J., Ganz, K., Steine, E.J., Cassady, J.P., Creyghton, M.P., Welstead, G.G., Gao, Q., Jaenisch, R. (2011). Reprogramming factor stoichiometry influences the epigenetic state and biological properties of induced pluripotent stem cells. Cell Stem Cell 9, 588–598.

Castel, G., Meistermann, D., Bretin, B., Firmin, J., Blin, J., Loubersac, S., Bruneau, A., Chevolleau, S., Kilens, S., Chariau, C., Gaignerie, A., Francheteau, Q., Kagawa, H., Charpentier, E., Flippe, L., Francois-Campion, V., Haider, S., Dietrich, B., Knofler, M., Arima, T., Bourdon, J., Rivron, N., Masson, D., Fournier, T., Okae, H., Freour, T., David, L. (2020). Induction of Human Trophoblast Stem Cells from Somatic Cells and Pluripotent Stem Cells. Cell Rep 33, 108419.

Chen, D., Liu, W., Zimmerman, J., Pastor, W.A., Kim, R., Hosohama, L., Ho, J., Aslanyan, M., Gell, J.J., Jacobsen, S.E., Clark, A.T. (2018). The TFAP2C-Regulated OCT4 Naive Enhancer Is Involved in Human Germline Formation. Cell Rep 25, 3591–3602 e3595.

Chiu, Y.H., Chen, H. (2016). GATA3 inhibits GCM1 activity and trophoblast cell invasion. Sci Rep 6, 21630.

Cinkornpumin, J.K., Kwon, S.Y., Guo, Y., Hossain, I., Sirois, J., Russett, C.S., Tseng, H.W., Okae, H., Arima, T., Duchaine, T.F., Liu, W., Pastor, W.A. (2020). Naive Human Embryonic Stem Cells Can Give Rise to Cells with a Trophoblast-like Transcriptome and Methylome. Stem Cell Reports 15, 198–213.

Costa, M.A. (2016). The endocrine function of human placenta: an overview. Reprod Biomed Online 32, 14–43.

Dong, C., Beltcheva, M., Gontarz, P., Zhang, B., Popli, P., Fischer, L.A., Khan, S.A., Park, K.M., Yoon, E.J., Xing, X., Kommagani, R., Wang, T., Solnica-Krezel, L., Theunissen, T.W. (2020). Derivation of trophoblast stem cells from naive human pluripotent stem cells. Elife 9.

Ellis, J. (2005). Silencing and variegation of gammaretrovirus and lentivirus vectors. Hum Gene Ther 16, 1241–1246.

Eloranta, J.J., Hurst, H.C. (2002). Transcription factor AP-2 interacts with the SUMO-conjugating enzyme UBC9 and is sumolated in vivo. J Biol Chem 277, 30798–30804.

Fabregat, A., Jupe, S., Matthews, L., Sidiropoulos, K., Gillespie, M., Garapati, P., Haw, R., Jassal, B., Korninger, F., May, B., Milacic, M., Roca, C.D., Rothfels, K., Sevilla, C., Shamovsky, V., Shorser, S., Varusai, T., Viteri, G., Weiser, J., Wu, G., Stein, L., Hermjakob, H., D’Eustachio, P. (2018). The Reactome Pathway Knowledgebase. Nucleic Acids Res D649-D655.

Fisher, S.J. (2015). Why is placentation abnormal in preeclampsia? Am J Obstet Gyneco l213, S115–122.

Fujiwara, Y., Chang, A.N., Williams, A.M., Orkin, S.H. (2004). Functional overlap of GATA-1 and GATA-2 in primitive hematopoietic development. Blood 103, 583–585.

Gafni, O., Weinberger, L., Mansour, A.A., Manor, Y.S., Chomsky, E., Ben-Yosef, D., Kalma, Y., Viukov, S., Maza, I., Zviran, A., Rais, Y., Shipony, Z., Mukamel, Z., Krupalnik, V., Zerbib, M., Geula, S., Caspi, I., Schneir, D., Shwartz, T., Gilad, S., Amann-Zalcenstein, D., Benjamin, S., Amit, I., Tanay, A., Massarwa, R., Novershtern, N., Hanna, J.H. (2013). Derivation of novel human ground state naive pluripotent stem cells. Nature 504, 282–286.

Gardner, D.K., Vella, P., Lane, M., Wagley, L., Schlenker, T., Schoolcraft, W.B. (1998). Culture and transfer of human blastocysts increases implantation rates and reduces the need for multiple embryo transfers. Fertil Steril 69, 84–88.

Genbacev, O., Krtolica, A., Kaelin, W., Fisher, S.J. (2001). Human cytotrophoblast expression of the von Hippel-Lindau protein is downregulated during uterine invasion in situ and upregulated by hypoxia in vitro. Dev Biol 233, 526–536.

Gene Ontology, C. (2021). The Gene Ontology resource: enriching a GOld mine. Nucleic Acids Res 49, D325–D334.

Gerri, C., McCarthy, A., Alanis-Lobato, G., Demtschenko, A., Bruneau, A., Loubersac, S., Fogarty, N.M.E., Hampshire, D., Elder, K., Snell, P., Christie, L., David, L., Van de Velde, H., Fouladi-Nashta, A.A., Niakan, K.K. (2020). Initiation of a conserved trophectoderm program in human, cow and mouse embryos. Nature 587, 443–447.

Guo, G., Stirparo, G.G., Strawbridge, S.E., Spindlow, D., Yang, J., Clarke, J., Dattani, A., Yanagida, A., Li, M.A., Myers, S., Ozel, B.N., Nichols, J., Smith, A. (2021). Human naive epiblast cells possess unrestricted lineage potential. Cell Stem Cell 28, 1040–1056 e1046.

Haider, S., Meinhardt, G., Saleh, L., Kunihs, V., Gamperl, M., Kaindl, U., Ellinger, A., Burkard, T.R., Fiala, C., Pollheimer, J., Mendjan, S., Latos, P.A., Knofler, M. (2018). Self-Renewing Trophoblast Organoids Recapitulate the Developmental Program of the Early Human Placenta. Stem Cell Reports 11, 537-551.

Hemberger, M., Hanna, C.W., Dean, W. (2020). Mechanisms of early placental development in mouse and humans. Nat Rev Genet 21, 27–43.

Herbst, F., Ball, C.R., Tuorto, F., Nowrouzi, A., Wang, W., Zavidij, O., Dieter, S.M., Fessler, S., van der Hoeven, F., Kloz, U., Lyko, F., Schmidt, M., von Kalle, C., Glimm, H. (2012). Extensive methylation of promoter sequences silences lentiviral transgene expression during stem cell differentiation in vivo. Mol Ther 20, 1014–1021.

Hertig, A.T., Rock, J., Adams, E (1956). A description of 34 human ova within the first 17 days of development. Am J Anat 98, 435–493.

Home, P., Kumar, R.P., Ganguly, A., Saha, B., Milano-Foster, J., Bhattacharya, B., Ray, S., Gunewardena, S., Paul, A., Camper, S.A., Fields, P.E., Paul, S. (2017). Genetic redundancy of GATA factors in the extraembryonic trophoblast lineage ensures the progression of preimplantation and postimplantation mammalian development. Development 144, 876–888.

Houghton, F.D. (2006). Energy metabolism of the inner cell mass and trophectoderm of the mouse blastocyst. Differentiation 74, 11–18.

Hu, H., Miao, Y.R., Jia, L.H., Yu, Q.Y., Zhang, Q., Guo, A.Y. (2019). AnimalTFDB 3.0: a comprehensive resource for annotation and prediction of animal transcription factors. Nucleic Acids Res 47, D33–D38.

Hyslop, L.A., Blakeley, P., Craven, L., Richardson, J., Fogarty, N.M., Fragouli, E., Lamb, M., Wamaitha, S.E., Prathalingam, N., Zhang, Q., O’Keefe, H., Takeda, Y., Arizzi, L., Alfarawati, S., Tuppen, H.A., Irving, L., Kalleas, D., Choudhary, M., Wells, D., Murdoch, A.P., Turnbull, D.M., Niakan, K.K., Herbert, M. (2016). Towards clinical application of pronuclear transfer to prevent mitochondrial DNA disease. Nature 534, 383–386.

Io, S., Kabata, M., Iemura, Y., Semi, K., Morone, N., Minagawa, A., Wang, B., Okamoto, I., Nakamura, T., Kojima, Y., Iwatani, C., Tsuchiya, H., Kaswandy, B., Kondoh, E., Kaneko, S., Woltjen, K., Saitou, M., Yamamoto, T., Mandai, M., Takashima, Y. (2021). Capturing human trophoblast development with naive pluripotent stem cells in vitro. Cell Stem Cell 28, 1023–1039 e1013.

Jassal, B., Matthews, L., Viteri, G., Gong, C., Lorente, P., Fabregat, A., Sidiropoulos, K., Cook, J., Gillespie, M., Haw, R., Loney, F., May, B., Milacic, M., Rothfels, K., Sevilla, C., Shamovsky, V., Shorser, S., Varusai, T., Weiser, J., Wu, G., Stein, L., Hermjakob, H., D’Eustachio, P. (2020). The reactome pathway knowledgebase. Nucleic Acids Res 48, D498–D503.

Jokimaa, V., Inki, P., Kujari, H., Hirvonen, O., Ekholm, E., Anttila, L. (1998). Expression of syndecan-1 in human placenta and decidua. Placenta 19, 157–163.

Kharchenko, P.V., Silberstein, L., Scadden, D.T. (2014). Bayesian approach to single-cell differential expression analysis. Nat Methods 11, 740–742.

Kidder, B.L., Palmer, S. (2010). Examination of transcriptional networks reveals an important role for TCFAP2C, SMARCA4, and EOMES in trophoblast stem cell maintenance. Genome Res 20, 458-472.

Kim, D., Paggi, J.M., Park, C., Bennett, C., Salzberg, S.L. (2019). Graph-based genome alignment and genotyping with HISAT2 and HISAT-genotype. Nat Biotechnol 37, 907–915.

Kliman, H.J., Nestler, J.E., Sermasi, E., Sanger, J.M., Strauss, J.F., 3rd (1986). Purification, characterization, and in vitro differentiation of cytotrophoblasts from human term placentae. Endocrinology 118, 1567-1582.

Kobayashi, N., Okae, H., Hiura, H., Kubota, N., Kobayashi, E.H., Shibata, S., Oike, A., Hori, T., Kikutake, C., Hamada, H., Kaji, H., Suyama, M., Bortolin-Cavaille, M.L., Cavaille, J., Arima, T. (2022). The microRNA cluster C19MC confers differentiation potential into trophoblast lineages upon human pluripotent stem cells. Nat Commun 13, 3071.

Kohama, K., Kawaguchi, M., Fukushima, T., Lin, C.Y., Kataoka, H. (2012). Regulation of pericellular proteolysis by hepatocyte growth factor activator inhibitor type 1 (HAI-1) in trophoblast cells. Hum Cell 25, 100–110.

Krendl, C., Shaposhnikov, D., Rishko, V., Ori, C., Ziegenhain, C., Sass, S., Simon, L., Muller, N.S., Straub, T., Brooks, K.E., Chavez, S.L., Enard, W., Theis, F.J., Drukker, M. (2017). GATA2/3-TFAP2A/C transcription factor network couples human pluripotent stem cell differentiation to trophectoderm with repression of pluripotency. Proc Natl Acad Sci U S A 114, E9579–E9588.

Kubaczka, C., Senner, C.E., Cierlitza, M., Arauzo-Bravo, M.J., Kuckenberg, P., Peitz, M., Hemberger, M., Schorle, H. (2015). Direct Induction of Trophoblast Stem Cells from Murine Fibroblasts. Cell Stem Cell 17, 557–568.

Kuckenberg, P., Buhl, S., Woynecki, T., van Furden, B., Tolkunova, E., Seiffe, F., Moser, M., Tomilin, A., Winterhager, E., Schorle, H. (2010). The transcription factor TCFAP2C/AP-2gamma cooperates with CDX2 to maintain trophectoderm formation. Mol Cell Biol 30, 3310–3320.

Law, C.W., Alhamdoosh, M., Su, S., Dong, X., Tian, L., Smyth, G.K., Ritchie, M.E. (2016). RNA-seq analysis is easy as 1-2-3 with limma, Glimma and edgeR. F1000Res 5.

Lee, C.Q., Gardner, L., Turco, M., Zhao, N., Murray, M.J., Coleman, N., Rossant, J., Hemberger, M., Moffett, A. (2016). What Is Trophoblast? A Combination of Criteria Define Human First-Trimester Trophoblast. Stem Cell Reports 6, 257–272.

Leese, H.J., Baumann, C.G., Brison, D.R., McEvoy, T.G., Sturmey, R.G. (2008). Metabolism of the viable mammalian embryo: quietness revisited. Mol Hum Reprod 14, 667–672.

Liao, Y., Smyth, G.K., Shi, W. (2019). The R package Rsubread is easier, faster, cheaper and better for alignment and quantification of RNA sequencing reads. Nucleic Acids Res 47, e47.

Lin, S.C., Wani, M.A., Whitsett, J.A., Wells, J.M. (2010). Klf5 regulates lineage formation in the pre-implantation mouse embryo. Development 137, 3953-3963.

Liu, X., Ouyang, J.F., Rossello, F.J., Tan, J.P., Davidson, K.C., Valdes, D.S., Schroder, J., Sun, Y.B.Y., Chen, J., Knaupp, A.S., Sun, G., Chy, H.S., Huang, Z., Pflueger, J., Firas, J., Tano, V., Buckberry, S., Paynter, J.M., Larcombe, M.R., Poppe, D., Choo, X.Y., O’Brien, C.M., Pastor, W.A., Chen, D., Leichter, A.L., Naeem, H., Tripathi, P., Das, P.P., Grubman, A., Powell, D.R., Laslett, A.L., David, L., Nilsson, S.K., Clark, A.T., Lister, R., Nefzger, C.M., Martelotto, L.G., Rackham, O.J.L., Polo, J.M. (2020). Reprogramming roadmap reveals route to human induced trophoblast stem cells. Nature 586, 101–107.

Love, M.I., Huber, W. (2014). Moderated estimation of fold change and dispersion for RNA-seq data with DESeq2. Genome Biol 15, 550.

Ma, G.T., Roth, M.E., Groskopf, J.C., Tsai, F.Y., Orkin, S.H., Grosveld, F., Engel, J.D., Linzer, D.I. (1997). GATA-2 and GATA-3 regulate trophoblast-specific gene expression in vivo. Development 124, 907–914.

Mandal, P.K., Rossi, D.J. (2013). Reprogramming human fibroblasts to pluripotency using modified mRNA. Nat Protoc 8, 568–582.

Meinhardt, G., Haider, S., Haslinger, P., Proestling, K., Fiala, C., Pollheimer, J., K nofler, M. (2014). Wnt-dependent T-cell factor-4 controls human etravillous trophoblast motility. Endocrinology 155, 1908–1920.

Meistermann, D., Bruneau, A., Loubersac, S., Reignier, A., Firmin, J., Francois-Campion, V., Kilens, S., Lelievre, Y., Lammers, J., Feyeux, M., Hulin, P., Nedellec, S., Bretin, B., Castel, G., Allegre, N., Covin, S., Bihouee, A., Soumillon, M., Mikkelsen, T., Barriere, P., Chazaud, C., Chappell, J., Pasque, V., Bourdon, J., Freour, T., David, L. (2021). Integrated pseudotime analysis of human pre-implantation embryo single-cell transcriptomes reveals the dynamics of lineage specification. Cell Stem Cell 28, 1625–1640 e1626.

Mi, H., Muruganujan, A., Ebert, D., Huang, X., Thomas, P.D. (2019). PANTHER version 14: more genomes, a new PANTHER GO-slim and improvements in enrichment analysis tools. Nucleic Acids Res 47, D419–D426.

Muraoka, N., Ied a, (2015). Stoichiometry of transcription factors is critical for cardiac reprogramming. Circ Res 116, 216–218.

Murthi, P., Kalionis, B., Cocquebert, M., Rajaraman, G., Chui, A., Keogh, R.J., Evain-Brion, D., Fournier, T. (2013). Homeobox genes and down-stream transcription factor PPARgamma in normal and pathological human placental development. Placenta 34, 299–309.

Naama, M., Rahamim, M., Zayat, V., Sebban, S., Radwan, A., Orzech, D., Lasry, R., Ifrah, A., Jaber, M., Sabag, O., Yassen, H., Khatib, A., Epsztejn-Litman, S., Novoselsky-Persky, M., Makedonski, K., Deri, N., Goldman-Wohl, D., Cedar, H., Yagel, S., Eiges, R., Buganim, Y. (2023). Pluripotency-independent induction of human trophoblast stem cells from fibroblasts. Nat Commun 14, 3359.

Nakagawa, M., Koyanagi, M., Tanabe, K., Takahashi, K., Ichisaka, T., Aoi, T., Okita, K., Mochiduki, Y., Takizawa, N., Yamanaka, S. (2008). Generation of induced pluripotent stem cells without Myc from mouse and human fibroblasts. Nat Biotechno l26, 101–106.

Nakagawa, M., Takizawa, N., Narita, M., Ichisaka, T., Yamanaka, S. (2010). Promotion of direct reprogramming by transformation-deficient Myc. Proc Natl Acad Sci USA 107, 14152-14157.

Niwa, H., Toyooka, Y., Shimosato, D., Strumpf, D., Takahashi, K., Yagi, R., Rossant, J. (2005). Interaction between Oct3/4 and Cdx2 determines trophectoderm differentiation. Cell 123, 917–929.

Ohgushi, M., Taniyama, N., Vandenbon, A., Eiraku, M. (2022). Delamination of trophoblast-like syncytia from the amniotic ectodermal analogue in human primed embryonic stem cell-based differentiation model. Cell Rep 39, 110973.

Okae, H., Toh, H., Sato, T., Hiura, H., Takahashi, S., Shirane, K., Kabayama, Y., Suyama, M., Sasaki, H., Arima, T. (2018). Derivation of Human Trophoblast Stem Cells. Cell Stem Cell 22, 50–63 e56.

Pastor, W.A., Liu, W., Chen, D., Ho, J., Kim, R., Hunt, T.J., Lukianchikov, A., Liu, X., Polo, J.M., Jacobsen, S.E., Clark, A.T. (2018). TFAP2C regulates transcription in human naive pluripotency by opening enhancers. Nat Cell Biol 20, 553–564.

Pelekanos, R.A., Sardesai, V.S., Futrega, K., Lott, W.B., Kuhn, M., Doran, M.R. (2016). Isolation and Expansion of Mesenchymal Stem/Stromal Cells Derived from Human Placenta Tissue. J Vis Exp.

Peterkin, T., Gibson, A., Loose, M., Patient, R. (2005). The roles of GATA-4, −5 and −6 in vertebrate heart development. Semin Cell Dev Biol 16, 83–94.

Pijnenborg, R., Dixon, G., Robertson, W.B., Brosens, I. (1980). Trophoblastic invasion of human decidua from 8 to 18 weeks of pregnancy. Placenta 1, 3–19.

Pirooznia, M., Nagarajan, V., Deng, Y. (2007). GeneVenn - A web application for comparing gene lists using Venn diagrams. Bioinformation 1, 420–422.

Ralston, A., Cox, B.J., Nishioka, N., Sasaki, H., Chea, E., Rugg-Gunn, P., Guo, G., Robson, P., Draper, J.S., Rossant, J. (2010). Gata3 regulates trophoblast development downstream of Tead4 and in parallel to Cdx2. Development 137, 395–403.

Richardson, B.D., Cheng, Y.H., Langland, R.A., Handwerger, S. (2001). Differential expression of AP-2gamma and AP-2alpha during human trophoblast differentiation. Life Sci 69, 2157–2165.

Riley, P., Anson-Cartwright, L., Cross, J.C. (1998). The Hand1 bHLH transcription factor is essential for placentation and cardiac morphogenesis. Nat Genet 18, 271–275.

Roberts, R.M., Ezashi, T., Sheridan, M.A., Yang, Y. (2018). Specification of trophoblast from embryonic stem cells exposed to BMP4. Biol Reprod 99, 212–224.

Russ, A.P., Wattler, S., Colledge, W.H., Aparicio, S.A., Carlton, M.B., Pearce, J.J., Barton, S.C., Surani, M.A., Ryan, K., Nehls, M.C., Wilson, V., Evans, M.J. (2000). Eomesodermin is required for mouse trophoblast development and mesoderm formation. Nature 404, 95–99.

Soncin, F., Khater, M., To, C., Pizzo, D., Farah, O., Wakeland, A., Arul Nambi Rajan, K., Nelson, K.K., Chang, C.W., Moretto-Zita, M., Natale, D.R., Laurent, L.C., Parast, M.M. (2018). Comparative analysis of mouse and human placentae across gestation reveals species-specific regulators of placental development. Development 145.

Strumpf, D., Mao, C.A., Yamanaka, Y., Ralston, A., Chawengsaks ophak, K., Beck, F., Rossant, J. (2005). Cdx2 is required for correct cell fate specification and differentiation of trophectoderm in the mouse blastocyst. Development 132, 2093–2102.

Takahashi, K., Tanabe, K., Ohnuki, M., Narita, M., Ichisaka, T., Tomoda, K., Yamanaka, S. (2007). Induction of pluripotent stem cells from adult human fibroblasts by defined factors. Cell 131, 861–872.

Takashima, Y., Guo, G., Loos, R., Nichols, J., Ficz, G., Krueger, F., Oxley, D., Santos, F., Clarke, J., Mansfield, W., Reik, W., Bertone, P., Smith, A. (2014). Resetting transcription factor control circuitry toward ground-state pluripotency in human. Cell 158, 1254–1269.

Tanaka, S., Kunath, T., Hadjantonakis, A.K., Nagy, A., Rossant, J. (1998). Promotion of trophoblast stem cell proliferation by FGF4. Science 282, 2072-2075.

Theunissen, T.W., Powell, B.E., Wang, H., Mitalipova, M., Faddah, D.A., Reddy, J., Fan, Z.P., Maetzel, D., Ganz, K., Shi, L., Lungjangwa, T., Imsoonthornruksa, S., Stelzer, Y., Rangarajan, S., D’Alessio, A., Zhang, J., Gao, Q., Dawlaty, M.M., Young, R.A., Gray, N.S., Jaenisch, R. (2014). Systematic Identification of Culture Conditions for Induction and Maintenance of Naive Human Pluripotency. Cell Stem Cell 15, 524–526.

Turco, M.Y., Gardner, L., Kay, R.G., Hamilton, R.S., Prater, M., Hollinshead, M.S., McWhinnie, A., Esposito, L., Fernando, R., Skelton, H., Reimann, F., Gribble, F.M., Sharkey, A., Marsh, S.G.E., O’Rahilly, S., Hemberger, M., Burton, G.J., Moffett, A. (2018). Trophoblast organoids as a model for maternal-fetal interactions during human placentation. Nature 564, 263–267.

Wamaitha, S.E., del Valle, I., Cho, L.T., Wei, Y., Fogarty, N.M., Blakeley, P., Sherwood, R.I., Ji, H., Niakan, K.K. (2015). Gata6 potently initiates reprograming of pluripotent and differentiated cells to extraembryonic endoderm stem cells. Genes Dev 29, 1239–1255.

Wamaitha, S.E., Grybel, K.J., Alanis-Lobato, G., Gerri, C., Ogushi, S., McCarthy, A., Mahadevaiah, S.K., Healy, L., Lea, R.A., Molina-Arcas, M., Devito, L.G., Elder, K., Snell, P., Christie, L., Downward, J., Turner, J.M.A., Niakan, K.K. (2020). IGF1-mediated human embryonic stem cell self-renewal recapitulates the embryonic niche. Nat Commun 11, 764.

Wang, L.J., Chen, C.P., Lee, Y.S., Ng, P.S., Chang, G.D., Pao, Y.H., Lo, H.F., Peng, C.H., Cheong, M.L., Chen, H. (2022). Functional antagonism between DeltaNp63alpha and GCM1 regulates human trophoblast stemness and differentiation. Nat Commun 13, 1626.

Warren, L., Manos, P.D., Ahfeldt, T., Loh, Y.H., Li, H., Lau, F., Ebina, W., Mandal, P.K., Smith, Z.D., Meissner, A., Daley, G.Q., Brack, A.S., Collins, J.J., Cowan, C., Schlaeger, T.M., Rossi, D.J. (2010). Highly efficient reprogramming to pluripotency and directed differentiation of human cells with synthetic modified mRNA. Cell Stem Cell 7, 618–630.

Werling, U., Schorle, H. (2002). Transcription factor gene AP-2 gamma essential for early murine development. Mol Cell Biol 22, 3149-3156.

Wernig, M., Lengner, C.J., Hanna, J., Lodato, M.A., Steine, E., Foreman, R., Staerk, J., Markoulaki, S., Jaenisch, R. (2008). A drug-inducible transgenic system for direct reprogramming of multiple somatic cell types. Nat Biotechnol 26, 916–924.

Xu, R.H., Chen, X., Li, D.S., Li, R., Addicks, G.C., Glennon, C., Zwaka, T.P., Thomson, J.A. (2002). BMP4 initiates human embryonic stem cell differentiation to trophoblast. Nat Biotechnol 20, 1261–1264.

Yu, J., Chia, J., Canning, C.A., Jones, C.M., Bard, F.A., Virshup, D.M. (2014). WLS retrograde transport to the endoplasmic reticulum during Wnt secretion. Dev Cell 29, 277–291.

