## supplementary figures for "Transcription factor-based transdifferentiation of human embryonic to trophoblast stem cells"

### Supplementary Figure 1

A

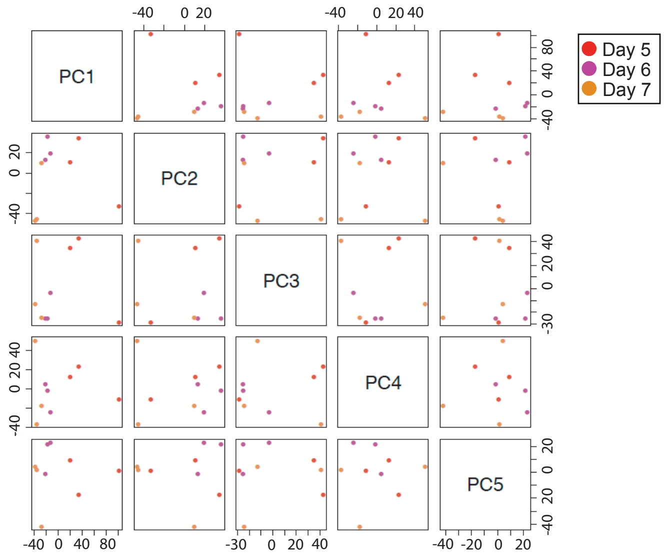


C

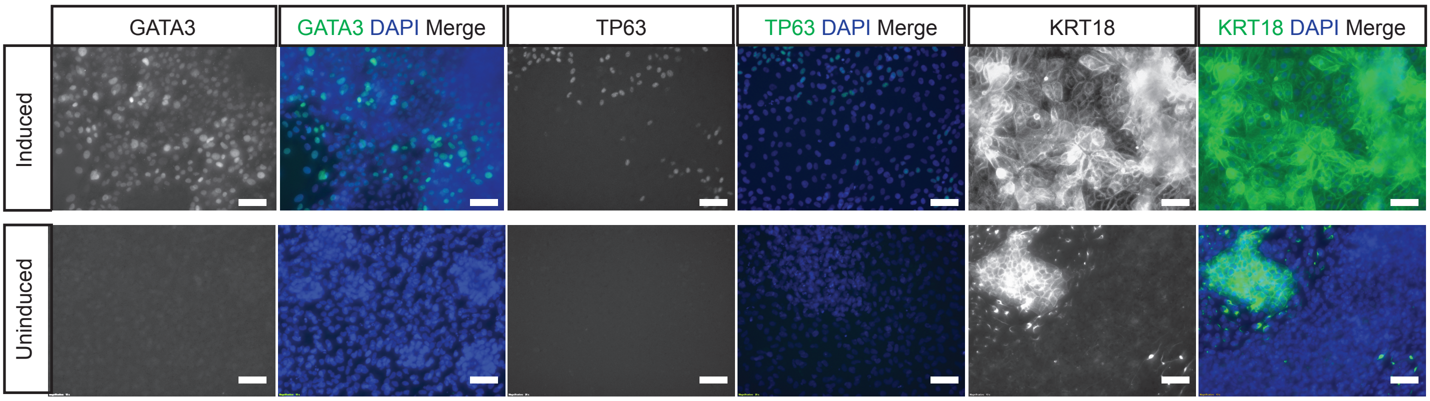

#### Supplementary Figure 2

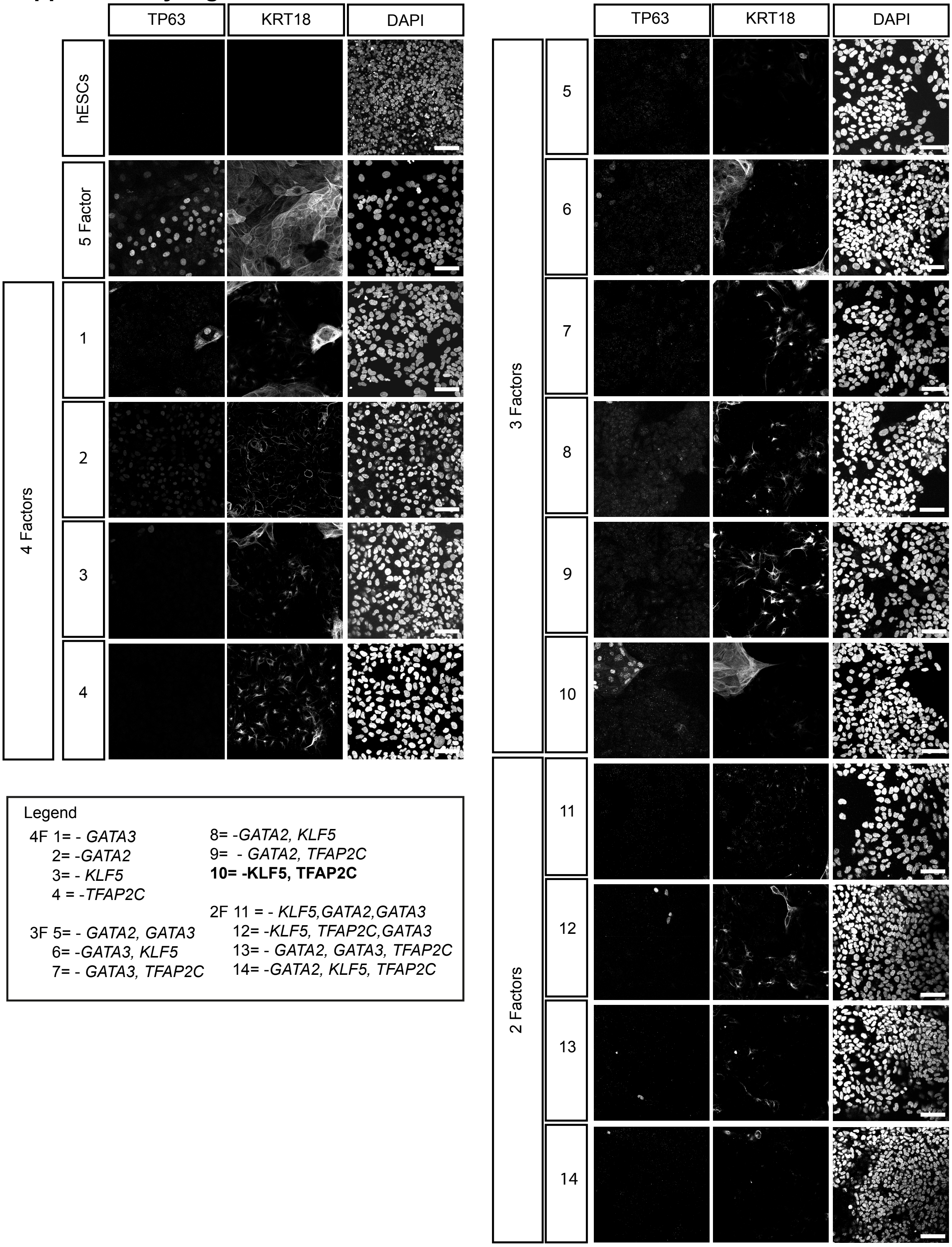

### Supplementary Figure 3

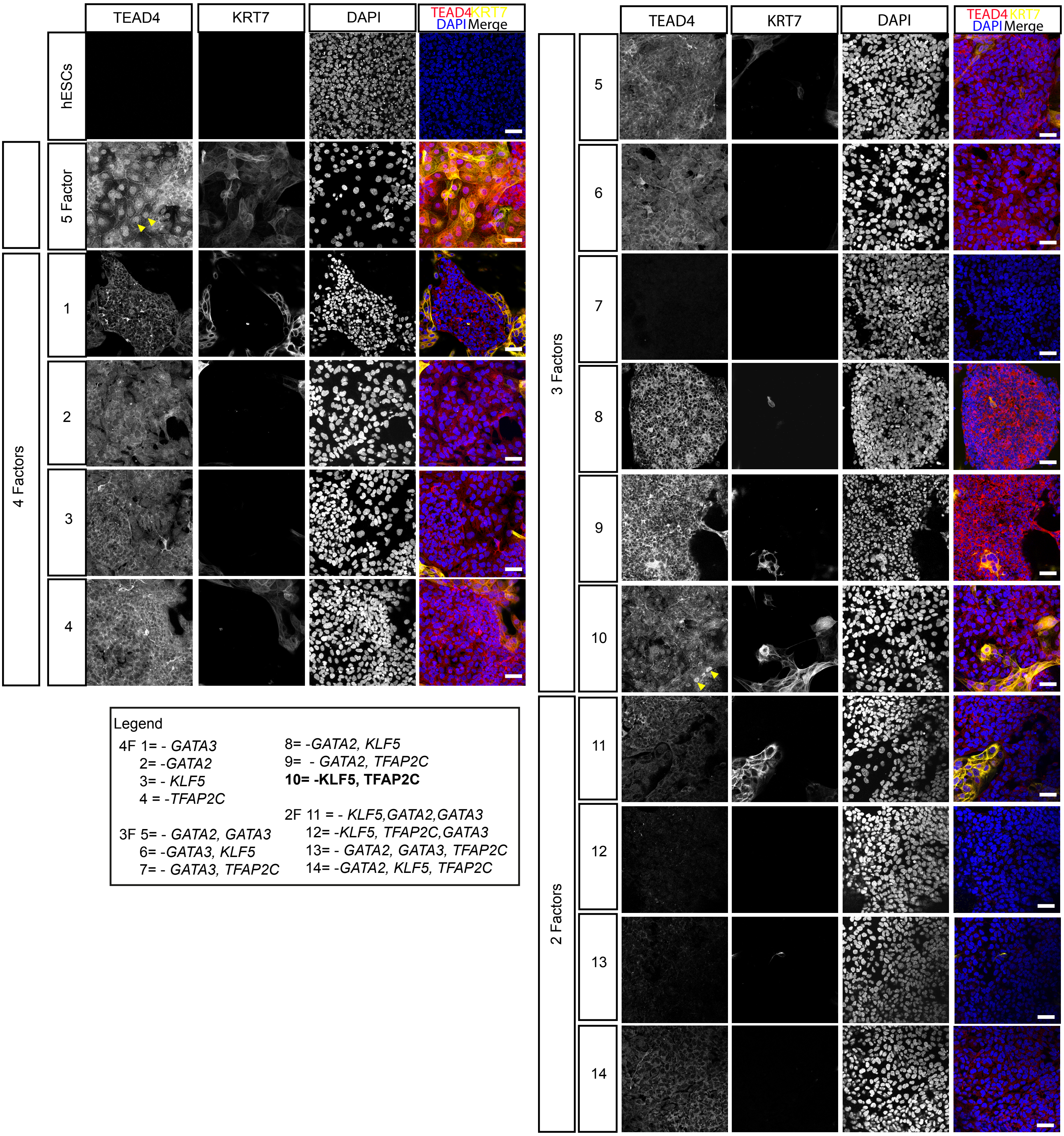


Legend

4F 1= - *GATA3*
2= -*GATA2*
3= - *KLF5*
4 = -*TFAP2C*

3F 5= - *GATA2, GATA3*
6= -*GATA3, KLF5*
7= - *GATA3, TFAP2C*
8= -*GATA2, KLF5*
9= - *GATA2, TFAP2C*
**10= -KLF5, TFAP2C**

2F 11 = - *KLF5,GATA2,GATA3*
12= -*KLF5, TFAP2C,GATA3*
13= - *GATA2, GATA3, TFAP2C*
14= -*GATA2, KLF5, TFAP2C*

**Figure S1.**

(A) Principal component analysis matrix using the first five principal components for the top 12,000 most variable expressed genes in day 5, 6 and 7 TE samples. (B) Immunofluorescence analysis for the detection of KRT18, GATA3 and TP63 (green) and DAPI nuclear expression (blue) in induced 5F-hESCs and uninduced 5F-hESCs in hTSC media on day 20. Scale bars: 50 µm.

**Figure S2 - GATA2 and GATA3 are required for induction of hTSC programme**

Extended immunofluorescence analysis for the detection of TP63, KRT18 and DAPI nuclear expression in transfected cells. Key for mRNA cocktail combinations: 1, *GATA2*, *TFAP2C*, *KLF5* and *MYC* (*GATA3* omitted); 2, *GATA3*, *TFAP2C*, *KLF5* and *MYC* (*GATA2* omitted); 3, *GATA2*, *GATA3*, *TFAP2C* and *MYC* (*KLF5* omitted); 4, *GATA2*, *GATA3*, *KLF5* and *MYC* (*TFAP2C* omitted); 5, *TFAP2C*, *KLF5* and *MYC* (*GATA2* and *GATA3* omitted); 6, *GATA2*, *TFAP2C* and *MYC* (*GATA3* and *KLF5* omitted); 7, *GATA2*, *KLF5* and *MYC* (*GATA3* and *TFAP2C* omitted); 8, *GATA3*, *TFAP2C* and *MYC* (*GATA2* and *KLF5* omitted); 9, *GATA3*, *KLF5* and *MYC* (*GATA*2 and *TFAP2C* omitted); 10, *GATA2*, *GATA3* and *MYC* (*TFAP2C* and *KLF5* omitted); 11, *TFAP2C*, *KLF5* and *MYC* (*KLF5*, *GATA2* and *GATA3* omitted); 12, *GATA2* and *MYC* (*GATA3*, *TFAP2C* and *KLF5*); 13, *KLF5* and *MYC* (*GATA2*, *GATA3* and *TFAP2C* omitted) and 14, *GATA3* and *MYC* (*GATA2*, *KLF5* and *TFAP2C* omitted). hESCs transfected with 5 factors are included as a positive control. hESCs cultured in mTeSR1 media are included as a negative control. Scale bars: 50 µm.

**Figure S3. Immunofluorescence analysis of additional markers in combinatorial experiments**

Immunofluorescence analysis for the detection of TEAD4, KRT7 and DAPI nuclear expression in transfected cells. Key for mRNA cocktail combinations: A, Five factors positive controls; 1, *GATA2*, *TFAP2C*, *KLF5* and *MYC* (*GATA3* omitted); 2, *GATA3*, *TFAP2C*, *KLF5* and *MYC* (*GATA2* omitted); 3, *GATA2*, *GATA3*, *TFAP2C* and *MYC* (*KLF5* omitted); 4, *GATA2*, *GATA3*, *KLF5* and *MYC* (*TFAP2C* omitted); 5, *TFAP2C*, *KLF5* and *MYC* (*GATA2* and *GATA3* omitted); 6, *GATA2*, *TFAP2C* and *MYC* (*GATA3* and *KLF5* omitted); 7, *GATA2*, *KLF5* and *MYC* (*GATA3* and *TFAP2C* omitted); 8, *GATA3*, *TFAP2C* and *MYC* (*GATA2* and *KLF5* omitted); 9, *GATA3*, *KLF5* and *MYC* (*GATA2* and *TFAP2C* omitted); 10, *GATA2*, *GATA3* and *MYC* (*TFAP2C* and *KLF5* omitted); 11, *TFAP2C*, *KLF5* and *MYC* (*KLF5*, *GATA2* and *GATA3* omitted); 12, *GATA2* and *MYC* (*GA*TA3,

*TFAP2C* and *KLF5*); 13, *KLF5* and *MYC* (*GATA2*, *GATA3* and *TFAP2C* omitted) and 14, *GATA3* and *MYC* (*GATA2*, *KLF5* and *TFAP2C* omitted). hESCs transfected with 5 factors are included as a positive control. hESCs culture in in mTeSR1 media are included as a negative control.

**Table S4. DESeq2 analysis to calculate the probability of differential expression between human TE versus EPI. The log2-fold change difference**

| **DESeq2 analysis** | | | | |
|---|---|---|---|---|
| | **TE mean** | **EPI mean** | **Log2FC** | **p-value** |
| GATA2 | 4528.6 | 16.1 | 8.1 | 2.20E-19 |
| GATA3 | 8457.3 | 255.1 | 5 | 4.06E-13 |
| KLF5 | 4609.4 | 1931.3 | 2.4 | 0.08 |
| TFAP2C | 1525.7 | 1135.1 | 0.4 | 0.9 |
| MYC | 3830.4 | 1405.5 | 1.4 | 0.6 |

| **RPKM values at time-points analysed** | | | |
|---|---|---|---|
| | **Day 5** | **Day 6** | **Day 7** |
| GATA2 | 19.3 | 65.6 | 52.5 |
| GATA3 | 93.8 | 120.4 | 96.8 |
| KLF5 | 45.2 | 45.2 | 46.7 |
| TFAP2C | 644.2 | 98.8 | 393.7 |
| MYC | 32.9 | 50.9 | 39.5 |

**Table S5. Primary antibodies used in this study**

| Antibody | Host | Supplier | Catalogue Number |
|---|---|---|---|
| OCT4 | Mouse IgG2b | Santa Cruz | SC-5279 |
| NANOG | Goat | R&D | AF1997 |
| GATA3 | Goat | R&D | AF2605 |
| TFAP2C | Goat | R&D | AF5059 |
| GATA2 | Rabbit | Santa Cruz | SC-9008 |
| NANOG | Rabbit | Abcam | ab21624 |
| KLF5 | Rabbit | Abcam | ab137676 |
| CYTOKERATIN18 | Mouse IgG1 | Abcam | ab668 |
| TP63 | Mouse | Biocare | 3066 |
| GCM1 | Rabbit | Sigma | HPA011343 |
| HAND1 | Rabbit | Abcam | ab196622 |
| TBX3 | Goat | Santa Cruz | SC-17871 |
| SOX2 | Rat | eBioscience | 14-9811-82 |
| TEAD4 | Mouse IgG2a | Abcam | ab58310 |
| CYTOKERATIN7 | Mouse | Agilent Dako | GA61961-2 |
| Alexa Fluor 594 Phalloidin | | ThermoFischer | A12381 |

#### Table S6. qRT-PCR primer sequences

| Oligo ID | Sequence | Reference | Oligo ID | Sequence |
|---|---|---|---|---|
| *EGFR* _F | CTAAGATCCCGTCCATCGCC | Soncin et al. 2018 | *PSG3* _F | TCGTAAAGCGAGGTGATGGG |
| *EGFR* _R | GGAGCCCAGCACTTTGATCT | Soncin et al. 2018 | *PSG3* _R | AAGCTCACAGCCTCCATGTC |
| *ENDOU* _F | ACAGGGCAGACACCAACAAA | | *SDC1* _F | CTTCACACTCCCCACACAGA |
| *ENDOU* _R | AGGAGGTTGATGAAGGCTGC | | *SDC1* _F | GTATTCTCCCCCGAGGTTTC |
| *ENPEP* _F | AATTTATGTCCAGCCAGAGC | | *TP63* _F | CTGGAAAACAATGCCCAGA |
| *ENPEP* _R | GTGATGAGTCCCCAGTTCTC | | *TP63* _R | AGAGAGCATCGAAGGTGGAG |
| *GABRP* _F | GCCCTAACAGAGCCTCAACA | Rostovkaya et al. 2022 | *WNT6* _F | CAGCCCCTTGGTTATGGACC |
| *GABRP* _R | CCCTGGATGCACATCCTCTC | Rostovkaya et al. 2022 | *WNT6* _R | TCTCCCGAATGTCCTGTTGC |
| *GAPDH* _F | GATGACATCAAGAAGGTGGTG | | | |
| *GAPDH* _R | GTCTACATGGCAACTGTGAGG | | | |
| *GATA2* _3UTR_F | AGGCCACTGACCATGAAGAA | | | |
| *GATA2* _3UTR_R | CGACGTCCATCTGTTCCCTA | | | |
| *GATA2* _F | GACTACAGCAGCGGACTCTT | | | |
| *GATA2* _R | GTTGTCGTCTGACAATTTGC | | | |
| *GATA3* _3UTR_F | GGTGTCTGTGTTCCAACCAC | | | |
| *GATA3* _3UTR_R | GTGGCCAGTGAAAGGAAACA | | | |
| *GATA3* _F | CCGCCCTACTACGGAAACTC | | | |
| GATA3_R | TCTTGGAGAAGGGGCTGAGA | | | |
| *GCM1* _F | TGCTGTCTGCTTCTCCGTAA | | | |
| *GCM1* _R | CACCTATTCGACTCCCCTCA | | | |
| *HEY1* _F | GCTGGTACCCAGTGCTTTTGAG | Io et al. 2021 | | |
| *HEY1* _R | CAAGGGCGTGCGCGTCAAAGTA | Io et al. 2021 | | |
| *HLAG* _F | CCACCACCCTGTCTTTGACTAT | Io et al. 2021 | | |
| *HLAG* _R | ACGTCCTGGGTCTGGTCCT | Io et al. 2021 | | |
| *ISL1* _F | TCTCCGGATTTGGAATGGCA | Yang et al. 2021 | | |
| *ISL1* _R | CCTTGCACCGCTTGTTTTGA | Yang et al. 2021 | | |
| *ITGA6* _F | GGCGGTGTTATGTCCTGAGTC | | | |
| *ITGA6* _R | AATCGCCCATCACAAAAGCTC | | | |
| *KLF5* _3UTR_F | GGGCTCCCTCAAATGACAGA | | | |
| *KLF5* _3UTR_R | CCACCCCTTACCCATGTTGA | | | |
| *KLF5* _F | CCACCACCCTGCCAGTTAAC | Takeda et al. 2022 | | |
| *KLF5* _R | TAAACTTTTGTGCAACCAGGGTAA | Takeda et al. 2022 | | |
| *LRRC32* _F | GCTGCACAACACCAAGACAAA | Io et al. 2021 | | |
| *LRRC32* _R | GATCAAGGGTCTCAGTGTCTGG | Io et al. 2021 | | |
| *LVRN* _F | GGGAGGGACTCTTCCTCAAC | Io et al. 2021 | | |
| *LVRN* _R | GGGAAAACATACCTGGCAAA | Io et al. 2021 | | |
| *MMP2* _F | CCCTGTGTCTTCCCCTTCAC | | | |
| *MMP2* _R | ATCGTAGTTGGCTGTGGTCG | | | |
| *MYC* _3UTR_F | ACCCTTCGCTATCATGCCTT | | | |
| *MYC* _3UTR_R | TCTTGGGCATGTGGATGAGT | | | |
| *MYC* _F | CGTCCTCGGATTCTCTGCTC | | | |
| *MYC* _R | GCTGCGTAGTTGTGCTGATG | | | |
| *NOTUM* _F | TTTGGCTACAAGGTCTACCCG | Io et al. 2021 | | |
| *NOTUM* _R | TCAAACAGCCACTGCACCAC | Io et al. 2021 | | |
| *NR2F2* _F | GCCATAGTCCTGTTCACCTCA | Io et al. 2021 | | |
| *NR2F2* _R | AATCTCGTCGGCTGGTTG | Io et al. 2021 | | |

**Table S7 Annotated sequence files of individual *in vitro* transcription template constructs for *GATA2*, *GATA3*, *TFAP2C* and *KLF5***

Bold and underlined: T7 promoter and T7 terminator
Bold: 5'UTR and 3'UTR

GATA2 isoform 1
**TAATACGACTCACTATAG**GG**AAATAAGAGAGAAAAGAAGAGTAAGAAGAAATATAAGAGCCAC**
**C**ATGGAGGTGGCGCCCGAGCAGCCGCGCTGGATGGCGCACCCGGCCGTGCTGAATGCGCAGCACC
CCGACTCACACCACCCGGGCCTGGCGCACAACTACATGGAACCCGCGCAGCTGCTGCCTCCAGACG
AGGTGGACGTCTTCTTCAATCACCTCGACTCGCAGGGCAACCCCTACTATGCCAACCCCGCTCACGC
GCGGGCGCGCGTCTCCTACAGCCCCGCGCACGCCCGCCTGACCGGAGGCCAGATGTGCCGCCCACA
CTTGTTGCACAGCCCGGGTTTGCCCTGGCTGGACGGGGGCAAAGCAGCCCTCTCTGCCGCTGCGGC
CCACCACCACAACCCCTGGACCGTGAGCCCCTTCTCCAAGACGCCACTGCACCCCTCAGCTGCTGGA
GGCCCTGGAGGCCCACTCTCTGTGTACCCAGGGGCTGGGGGTGGGAGCGGGGGAGGCAGCGGGA
GCTCAGTGGCCTCCCTCACCCCTACAGCAGCCCACTCTGGCTCCCACCTTTTCGGCTTCCCACCCACG
CCACCCAAAGAAGTGTCTCCTGACCCTAGCACCACGGGGGCTGCGTCTCCAGCCTCATCTTCCGCGG
GGGGTAGTGCAGCCCGAGGAGAGGACAAGGACGGCGTCAAGTACCAGGTGTCACTGACGGAGAG
CATGAAGATGGAAAGTGGCAGTCCCCTGCGCCCAGGCCTAGCTACTATGGGCACCCAGCCTGCTAC
ACACCACCCCATCCCCACCTACCCCTCCTATGTGCCGGCGGCTGCCCACGACTACAGCAGCGGACTCT
TCCACCCCGGAGGCTTCCTGGGGGGACCGGCCTCCAGCTTCACCCCTAAGCAGCGCAGCAAGGCTC
GTTCCTGTTCAGAAGGCCGGGAGTGTGTCAACTGTGGGGCCACAGCCACCCCTCTCTGGCGGCGGG
ACGGCACCGGCCACTACCTGTGCAATGCCTGTGGCCTCTACCACAAGATGAATGGGCAGAACCGAC
CACTCATCAAGCCCAAGCGAAGACTGTCGGCCGCCAGAAGAGCCGGCACCTGTTGTGCAAATTGTC
AGACGACAACCACCACCTTATGGCGCCGAAACGCCAACGGGGACCCTGTCTGCAACGCCTGTGGCC
TCTACTACAAGCTGCACAATGTTAACAGGCCACTGACCATGAAGAAGGAAGGGATCCAGACTCGGA
ACCGGAAGATGTCCAACAAGTCCAAGAAGAGCAAGAAAGGGGCGGAGTGCTTCGAGGAGCTGTCA
AAGTGCATGCAGGAGAAGTCATCCCCCTTCAGTGCAGCTGCCCTGGCTGGACACATGGCACCTGTG
GGCCACCTCCCGCCCTTCAGCCACTCCGGACACATCCTGCCCACTCCGACGCCCATCCACCCCTCCTC
CAGCCTCTCCTTCGGCCACCCCCACCCGTCCAGCATGGTGACCGCCATGGGCTAG**GCTGCCTTCTGC**
**GGGGCTTGCCTTCTGGCCATGCCCTTCTTCTCTCCCTTGCACCTGTACCTCTTGGTCTTTGAATAAAG**
**CCTGAGTAGGAAGTAGCATAACCCCTTGGGGCCTCTAAACGGGTCTTGAGGGGTTTTTTG**

GATA3 isoform 1
**TAATACGACTCACTATAG**GG**AAATAAGAGAGAAAAGAAGAGTAAGAAGAAATATAAGAGCCA**C
CATGGAGGTGACGGCGGACCAGCCGCGCTGGGTGAGCCACCACCACCCCGCCGTGCTCAACGGGC
AGCACCCGGACACGCACCACCCGGGCCTCAGCCACTCCTACATGGACGCGGCGCAGTACCCGCTGC
CGGAGGAGGTGGATGTGCTTTTTAACATCGACGGTCAAGGCAACCACGTCCCGCCCTACTACGGAA
ACTCGGTCAGGGCCACGGTGCAGAGGTACCCTCCGACCCACCACGGGAGCCAGGTGTGCCGCCCGC
CTCTGCTTCATGGATCCCTACCCTGGCTGGACGGCGGCAAAGCCCTGGGCAGCCACCACACCGCCTC
CCCCTGGAATCTCAGCCCCTTCTCCAAGACGTCCATCCACCACGGCTCCCCGGGGCCCCTCTCCGTCT
ACCCCCCGGCCTCGTCCTCCTCCTTGTCGGGGGGCCACGCCAGCCCGCACCTCTTCACCTTCCCGCCC
ACCCCGCCGAAGGACGTCTCCCCGGACCCATCGCTGTCCACCCCAGGCTCGGCCGGCTCGGCCCGG
CAGGACGAGAAAGAGTGCCTCAAGTACCAGGTGCCCCTGCCCGACAGCATGAAGCTGGAGTCGTCC
CACTCCCGTGGCAGCATGACCGCCCTGGGTGGAGCCTCCTCGTCGACCCACCACCCCATCACCACCT
ACCCGCCCTACGTGCCCGAGTACAGCTCCGGACTCTTCCCCCCCAGCAGCCTGCTGGGCGGCTCCCC
CACCGGCTTCGGATGCAAGTCCAGGCCCAAGGCCCGGTCCAGCACAGAAGGCAGGGAGTGTGTGA

ACTGTGGGGCAACCTCGACCCCACTGTGGCGGCGAGATGGCACGGGACACTACCTGTGCAACGCCT
GCGGGCTCTATCACAAAATGAACGGACAGAACCGGCCCCTCATTAAGCCCAAGCGAAGGCTGTCTG
CAGCCAGGAGAGCAGGGACGTCCTGTGCGAACTGTCAGACCACCACAACCACACTCTGGAGGAGG
AATGCCAATGGGGACCCTGTCTGCAATGCCTGTGGGCTCTACTACAAGCTTCACAATATTAACAGAC
CCCTGACTATGAAGAAGGAAGGCATCCAGACCAGAAACCGAAAAATGTCTAGCAAATCCAAAAAGT
GCAAAAAAGTGCATGACTCACTGGAGGACTTCCCCAAGAACAGCTCGTTTAACCCGGCCGCCCTCTC
CAGACACATGTCCTCCCTGAGCCACATCTCGCCCTTCAGCCACTCCAGCCACATGCTGACCACGCCCA
CGCCGATGCACCCGCCATCCAGCCTGTCCTTTGGACCACACCACCCCTCCAGCATGGTCACCGCCATG
GGTTAG**GCTGCCTTCTGCGGGGCTTGCCTTCTGGCCATGCCCTTCTTCTCTCCCTTGCACCTGTACCT
CTTGGTCTTTGAATAAAGCCTGAGTAGGAAGTAGCATAACCCCTTGGGGCCTCTAAACGGGTCTTG
AGGGGTTTTTTG**

KLF5 isoform 1
**TAATACGACTCACTATAG**GG**AAATAAGAGAGAAAAGAAGAGTAAGAAGAAATATAAGAGCCAC
C**ATGGCTACAAGGGTGCTGAGCATGAGCGCCCGCCTGGGACCCGTGCCCCAGCCGCCGGCGCCGCA
GGACGAGCCGGTGTTCGCGCAGCTCAAGCCGGTGCTGGGCGCCGCGAATCCGGCCCGCGACGCGG
CGCTCTTCCCCGGCGAGGAGCTGAAGCACGCGCACCACCGCCCGCAGGCGCAGCCCGCGCCCGCGC
AGGCCCCGCAGCCGGCCCAGCCGCCCGCCACCGGCCCGCGGCTGCCTCCAGAGGACCTGGTCCAGA
CAAGATGTGAAATGGAGAAGTATCTGACACCTCAGCTTCCTCCAGTTCCTATAATTCCAGAGCATAA
AAAGTATAGACGAGACAGTGCCTCAGTCGTAGACCAGTTCTTCACTGACACTGAAGGGTTACCTTAC
AGTATCAACATGAACGTCTTCCTCCCTGACATCACTCACCTGAGAACTGGCCTCTACAAATCCCAGAG
ACCGTGCGTAACACACATCAAGACAGAACCTGTTGCCATTTTCAGCCACCAGAGTGAAACGACTGCC
CCTCCTCCGGCCCCGACCCAGGCCCTCCCTGAGTTCACCAGTATATTCAGCTCACACCAGACCGCAGC
TCCAGAGGTGAACAATATTTTCATCAAACAAGAACTTCCTACACCAGATCTTCATCTTTCTGTCCCTAC
CCAGCAGGGCCACCTGTACCAGCTACTGAATACACCGGATCTAGATATGCCCAGTTCTACAAATCAG
ACAGCAGCAATGGACACTCTTAATGTTTCTATGTCAGCTGCCATGGCAGGCCTTAACACACACACCTC
TGCTGTTCCGCAGACTGCAGTGAAACAATTCCAGGGCATGCCCCCTTGCACATACACAATGCCAAGT
CAGTTTCTTCCACAACAGGCCACTTACTTTCCCCCGTCACCACCAAGCTCAGAGCCTGGAAGTCCAGA
TAGACAAGCAGAGATGCTCCAGAATTTAACCCCACCTCCATCCTATGCTGCTACAATTGCTTCTAAAC
TGGCAATTCACAATCCAAATTTACCCACCACCCTGCCAGTTAACTCACAAAACATCCAACCTGTCAGA
TACAATAGAAGGAGTAACCCCGATTTGGAGAAACGACGCATCCACTACTGCGATTACCCTGGTTGCA
CAAAAGTTTATACCAAGTCTTCTCATTTAAAAGCTCACCTGAGGACTCACACTGGTGAAAAGCCATAC
AAGTGTACCTGGGAAGGCTGCGACTGGAGGTTCGCGCGATCGGATGAGCTGACCCGCCACTACCG
GAAGCACACAGGCGCCAAGCCCTTCCAGTGCGGGGTGTGCAACCGCAGCTTCTCGCGCTCTGACCA
CCTGGCCCTGCATATGAAGAGGCACCAGAACTGA**GCTGCCTTCTGCGGGGCTTGCCTTCTGGCCAT
GCCCTTCTTCTCTCCCTTGCACCTGTACCTCTTGGTCTTTGAATAAAGCCTGAGTAGGAAGTAGCAT
AACCCCTTGGGGCCTCTAAACGGGTCTTGAGGGGTTTTTTG**

MYC
**TAATACGACTCACTATAG**GG**AAATAAGAGAGAAAAGAAGAGTAAGAAGAAATATAAGAGCCAC
C**ATGCCCCTCAACGTTAGCTTCACCAACAGGAACTATGACCTCGACTACGACTCGGTGCAGCCGTAT
TTCTACTGCGACGAGGAGGAGAACTTCTACCAGCAGCAGCAGCAGAGCGAGCTGCAGCCCCCGGC
GCCCAGCGAGGATATCTGGAAGAAATTCGAGCTGCTGCCCACCCCGCCCCTGTCCCCTAGCCGCCGC
TCCGGGCTCTGCTCGCCCTCCTACGTTGCGGTCACACCCTTCTCCCTTCGGGGAGACAACGACGGCG
GTGGCGGGAGCTTCTCCACGGCCGACCAGCTGGAGATGGTGACCGAGCTGCTGGGAGGAGACATG
GTGAACCAGAGTTTCATCTGCGACCCGGACGACGAGACCTTCATCAAAAACATCATCATCCAGGACT
GTATGTGGAGCGGCTTCTCGGCCGCCGCCAAGCTCGTCTCAGAGAAGCTGGCCTCCTACCAGGCTG

CGCGCAAAGACAGCGGCAGCCCGAACCCCGCCCGCGGCCACAGCGTCTGCTCCACCTCCAGCTTGT
ACCTGCAGGATCTGAGCGCCGCCGCCTCAGAGTGCATCGACCCCTCGGTGGTCTTCCCCTACCCTCT
CAACGACAGCAGCTCGCCCAAGTCCTGCGCCTCGCAAGACTCCAGCGCCTTCTCTCCGTCCTCGGATT
CTCTGCTCTCCTCGACGGAGTCCTCCCCGCAGGGCAGCCCCGAGCCCCTGGTGCTCCATGAGGAGAC
ACCGCCCACCACCAGCAGCGACTCTGAGGAGGAACAAGAAGATGAGGAAGAAATCGATGTTGTTTC
TGTGGAAAAGAGGCAGGCTCCTGGCAAAAGGTCAGAGTCTGGATCACCTTCTGCTGGAGGCCACAG
CAAACCTCCTCACAGCCCACTGGTCCTCAAGAGGTGCCACGTCTCCACACATCAGCACAACTACGCA
GCGCCTCCCTCCACTCGGAAGGACTATCCTGCTGCCAAGAGGGTCAAGTTGGACAGTGTCAGAGTC
CTGAGACAGATCAGCAACAACCGAAAATGCACCAGCCCCAGGTCCTCGGACACCGAGGAGAATGTC
AAGAGGCGAACACACAACGTCTTGGAGCGCCAGAGGAGGAACGAGCTAAAACGGAGCTTTTTTGC
CCTGCGTGACCAGATCCCGGAGTTGGAAAACAATGAAAAGGCCCCCAAGGTAGTTATCCTTAAAAA
AGCCACAGCATACATCCTGTCCGTCCAAGCAGAGGAGCAAAAGCTCATTTCTGAAGAGGACTTGTTG
CGGAAACGACGAGAACAGTTGAAACACAAACTTGAACAGCTACGGAACTCTTGTGCGTAA**GCTGCC**
**TTCTGCGGGGCTTGCCTTCTGGCCATGCCCTTCTTCTCTCCCTTGCACCTGTACCTCTTGGTCTTTGAA**
**TAAAGCCTGAGTAGGAAG<u>TAGCATAACCCCTTGGGGCCTCTAAACGGGTCTTGAGGGGTTTTTTG</u>**

TFAP2C isoform 1
**<u>TAATACGACTCACTATAG</u>**GG**AAATAAGAGAGAAAAGAAGAGTAAGAAGAAATATAAGAGCCAC**
**C**ATGTTGTGGAAAATAACCGATAATGTCAAGTACGAAGAGGACTGCGAGGATCGCCACGACGGGA
GCAGCAATGGGAATCCGCGGGTCCCCCACCTCTCCTCCGCCGGGCAGCACCTCTACAGCCCCGCGCC
ACCCCTCTCCCACACTGGAGTCGCCGAATATCAGCCGCCACCCTACTTTCCCCCTCCCTACCAGCAGC
TGGCCTACTCCCAGTCGGCCGACCCCTACTCGCATCTGGGGGAAGCGTACGCCGCCGCCATCAACCC
CCTGCACCAGCCGGCGCCCACAGGCAGCCAGCAGCAGGCCTGGCCCGGCCGCCAGAGCCAGGAGG
GAGCGGGGCTGCCCTCGCACCACGGGCGCCCGGCCGGCCTACTGCCCCACCTCTCCGGGCTGGAGG
CGGGCGCGGTGAGCGCCCGCAGGGATGCCTACCGCCGCTCCGACCTGCTGCTGCCCCACGCACACG
CCCTGGATGCCGCGGGCCTGGCCGAGAACCTGGGGCTCCACGACATGCCTCACCAGATGGACGAG
GTGCAGAATGTCGACGACCAGCACCTGTTGCTGCACGATCAGACAGTCATTCGCAAAGGTCCCATTT
CCATGACCAAGAACCCTCTGAACCTCCCCTGTCAGAAGGAGCTGGTGGGGGCCGTAATGAACCCCA
CTGAGGTCTTCTGCTCAGTCCCTGGAAGATTGTCGCTCCTCAGCTCTACGTCTAAATACAAAGTGACA
GTGGCTGAAGTACAGAGGCGACTGTCCCCACCTGAATGCTTAAATGCCTCGTTACTGGGAGGTGTTC
TCAGAAGAGCCAAATCGAAAAATGGAGGCCGGTCCTTGCGGGAGAAGTTGGACAAGATTGGGTTG
AATCTTCCGGCCGGGAGGCGGAAAGCCGCTCATGTGACTCTCCTGACATCCTTAGTAGAAGGTGAA
GCTGTTCATTTGGCTAGGGACTTTGCCTATGTCTGTGAAGCCGAATTTCCTAGTAAACCAGTGGCAG
AATATTTAACCAGACCTCATCTTGGAGGACGAAATGAGATGGCAGCTAGGAAGAACATGCTATTGG
CGGCCCAGCAACTGTGTAAAGAATTCACAGAACTTCTCAGCCAAGACCGGACACCCCATGGACCA
GCAGGCTCGCCCCAGTCTTGGAGACGAACATACAGAACTGCTTGTCTCATTTCAGCCTGATTACCCA
CGGGTTTGGCAGCCAGGCCATCTGTGCCGCGGTGTCTGCCCTGCAGAACTACATCAAAGAAGCCCT
GATTGTCATAGACAAATCCTACATGAACCCTGGAGACCAGAGTCCAGCTGATTCTAACAAAACCCTG
GAGAAAATGGAGAAACACAGGAAATAA**GCTGCCTTCTGCGGGGCTTGCCTTCTGGCCATGCCCTTC**
**TTCTCTCCCTTGCACCTGTACCTCTTGGTCTTTGAATAAAGCCTGAGTAGGAAG<u>TAGCATAACCCCT</u>**
**<u>TGGGGCCTCTAAACGGGTCTTGAGGGGTTTTTTG</u>**
