## Supplementary table 7 for "Transcription factor-based transdifferentiation of human embryonic to trophoblast stem cells"

Bold and underlined: T7 promoter and T7 terminator

Bold: 5’UTR and 3’UTR

GATA2 isoform 1

**TAATACGACTCACTATAG**GG**AAATAAGAGAGAAAAGAAGAGTAAGAAGAAATATAAGAGCCACC**ATGGAGGTGGCGCCCGAGCAGCCGCGCTGGATGGCGCACCCGGCCGTGCTGAATGCGCAGCACCCCGACTCACACCACCCGGGCCTGGCGCACAACTACATGGAACCCGCGCAGCTGCTGCCTCCAGACGAGGTGGACGTCTTCTTCAATCACCTCGACTCGCAGGGCAACCCCTACTATGCCAACCCCGCTCACGCGCGGGCGCGCGTCTCCTACAGCCCCGCGCACGCCCGCCTGACCGGAGGCCAGATGTGCCGCCCACACTTGTTGCACAGCCCGGGTTTGCCCTGGCTGGACGGGGGCAAAGCAGCCCTCTCTGCCGCTGCGGCCCACCACCACAACCCCTGGACCGTGAGCCCCTTCTCCAAGACGCCACTGCACCCCTCAGCTGCTGGAGGCCCTGGAGGCCCACTCTCTGTGTACCCAGGGGCTGGGGGTGGGAGCGGGGGAGGCAGCGGGAGCTCAGTGGCCTCCCTCACCCCTACAGCAGCCCACTCTGGCTCCCACCTTTTCGGCTTCCCACCCACGCCACCCAAAGAAGTGTCTCCTGACCCTAGCACCACGGGGGCTGCGTCTCCAGCCTCATCTTCCGCGGGGGGTAGTGCAGCCCGAGGAGAGGACAAGGACGGCGTCAAGTACCAGGTGTCACTGACGGAGAGCATGAAGATGGAAAGTGGCAGTCCCCTGCGCCCAGGCCTAGCTACTATGGGCACCCAGCCTGCTACACACCACCCCATCCCCACCTACCCCTCCTATGTGCCGGCGGCTGCCCACGACTACAGCAGCGGACTCTTCCACCCCGGAGGCTTCCTGGGGGGACCGGCCTCCAGCTTCACCCCTAAGCAGCGCAGCAAGGCTCGTTCCTGTTCAGAAGGCCGGGAGTGTGTCAACTGTGGGGCCACAGCCACCCCTCTCTGGCGGCGGGACGGCACCGGCCACTACCTGTGCAATGCCTGTGGCCTCTACCACAAGATGAATGGGCAGAACCGACCACTCATCAAGCCCAAGCGAAGACTGTCGGCCGCCAGAAGAGCCGGCACCTGTTGTGCAAATTGTCAGACGACAACCACCACCTTATGGCGCCGAAACGCCAACGGGGACCCTGTCTGCAACGCCTGTGGCCTCTACTACAAGCTGCACAATGTTAACAGGCCACTGACCATGAAGAAGGAAGGGATCCAGACTCGGAACCGGAAGATGTCCAACAAGTCCAAGAAGAGCAAGAAAGGGGCGGAGTGCTTCGAGGAGCTGTCAAAGTGCATGCAGGAGAAGTCATCCCCCTTCAGTGCAGCTGCCCTGGCTGGACACATGGCACCTGTGGGCCACCTCCCGCCCTTCAGCCACTCCGGACACATCCTGCCCACTCCGACGCCCATCCACCCCTCCTCCAGCCTCTCCTTCGGCCACCCCCACCCGTCCAGCATGGTGACCGCCATGGGCTAG**GCTGCCTTCTGCGGGGCTTGCCTTCTGGCCATGCCCTTCTTCTCTCCCTTGCACCTGTACCTCTTGGTCTTTGAATAAAGCCTGAGTAGGAAGTAGCATAACCCCTTGGGGCCTCTAAACGGGTCTTGAGGGGTTTTTTG**

GATA3 isoform 1

**TAATACGACTCACTATAG**GG**AAATAAGAGAGAAAAGAAGAGTAAGAAGAAATATAAGAGCCACC**ATGGAGGTGACGGCGGACCAGCCGCGCTGGGTGAGCCACCACCACCCCGCCGTGCTCAACGGGCAGCACCCGGACACGCACCACCCGGGCCTCAGCCACTCCTACATGGACGCGGCGCAGTACCCGCTGCCGGAGGAGGTGGATGTGCTTTTTAACATCGACGGTCAAGGCAACCACGTCCCGCCCTACTACGGAAACTCGGTCAGGGCCACGGTGCAGAGGTACCCTCCGACCCACCACGGGAGCCAGGTGTGCCGCCCGCCTCTGCTTCATGGATCCCTACCCTGGCTGGACGGCGGCAAAGCCCTGGGCAGCCACCACACCGCCTCCCCCTGGAATCTCAGCCCCTTCTCCAAGACGTCCATCCACCACGGCTCCCCGGGGCCCCTCTCCGTCTACCCCCCGGCCTCGTCCTCCTCCTTGTCGGGGGGCCACGCCAGCCCGCACCTCTTCACCTTCCCGCCCACCCCGCCGAAGGACGTCTCCCCGGACCCATCGCTGTCCACCCCAGGCTCGGCCGGCTCGGCCCGGCAGGACGAGAAAGAGTGCCTCAAGTACCAGGTGCCCCTGCCCGACAGCATGAAGCTGGAGTCGTCCCACTCCCGTGGCAGCATGACCGCCCTGGGTGGAGCCTCCTCGTCGACCCACCACCCCATCACCACCTACCCGCCCTACGTGCCCGAGTACAGCTCCGGACTCTTCCCCCCCAGCAGCCTGCTGGGCGGCTCCCCCACCGGCTTCGGATGCAAGTCCAGGCCCAAGGCCCGGTCCAGCACAGAAGGCAGGGAGTGTGTGAACTGTGGGGCAACCTCGACCCCACTGTGGCGGCGAGATGGCACGGGACACTACCTGTGCAACGCCTGCGGGCTCTATCACAAAATGAACGGACAGAACCGGCCCCTCATTAAGCCCAAGCGAAGGCTGTCTGCAGCCAGGAGAGCAGGGACGTCCTGTGCGAACTGTCAGACCACCACAACCACACTCTGGAGGAGGAATGCCAATGGGGACCCTGTCTGCAATGCCTGTGGGCTCTACTACAAGCTTCACAATATTAACAGACCCCTGACTATGAAGAAGGAAGGCATCCAGACCAGAAACCGAAAAATGTCTAGCAAATCCAAAAAGTGCAAAAAAGTGCATGACTCACTGGAGGACTTCCCCAAGAACAGCTCGTTTAACCCGGCCGCCCTCTCCAGACACATGTCCTCCCTGAGCCACATCTCGCCCTTCAGCCACTCCAGCCACATGCTGACCACGCCCACGCCGATGCACCCGCCATCCAGCCTGTCCTTTGGACCACACCACCCCTCCAGCATGGTCACCGCCATGGGTTAG**GCTGCCTTCTGCGGGGCTTGCCTTCTGGCCATGCCCTTCTTCTCTCCCTTGCACCTGTACCTCTTGGTCTTTGAATAAAGCCTGAGTAGGAAGTAGCATAACCCCTTGGGGCCTCTAAACGGGTCTTGAGGGGTTTTTTG**

KLF5 isoform 1

**TAATACGACTCACTATAG**GG**AAATAAGAGAGAAAAGAAGAGTAAGAAGAAATATAAGAGCCACC**ATGGCTACAAGGGTGCTGAGCATGAGCGCCCGCCTGGGACCCGTGCCCCAGCCGCCGGCGCCGCAGGACGAGCCGGTGTTCGCGCAGCTCAAGCCGGTGCTGGGCGCCGCGAATCCGGCCCGCGACGCGGCGCTCTTCCCCGGCGAGGAGCTGAAGCACGCGCACCACCGCCCGCAGGCGCAGCCCGCGCCCGCGCAGGCCCCGCAGCCGGCCCAGCCGCCCGCCACCGGCCCGCGGCTGCCTCCAGAGGACCTGGTCCAGACAAGATGTGAAATGGAGAAGTATCTGACACCTCAGCTTCCTCCAGTTCCTATAATTCCAGAGCATAAAAAGTATAGACGAGACAGTGCCTCAGTCGTAGACCAGTTCTTCACTGACACTGAAGGGTTACCTTACAGTATCAACATGAACGTCTTCCTCCCTGACATCACTCACCTGAGAACTGGCCTCTACAAATCCCAGAGACCGTGCGTAACACACATCAAGACAGAACCTGTTGCCATTTTCAGCCACCAGAGTGAAACGACTGCCCCTCCTCCGGCCCCGACCCAGGCCCTCCCTGAGTTCACCAGTATATTCAGCTCACACCAGACCGCAGCTCCAGAGGTGAACAATATTTTCATCAAACAAGAACTTCCTACACCAGATCTTCATCTTTCTGTCCCTACCCAGCAGGGCCACCTGTACCAGCTACTGAATACACCGGATCTAGATATGCCCAGTTCTACAAATCAGACAGCAGCAATGGACACTCTTAATGTTTCTATGTCAGCTGCCATGGCAGGCCTTAACACACACACCTCTGCTGTTCCGCAGACTGCAGTGAAACAATTCCAGGGCATGCCCCCTTGCACATACACAATGCCAAGTCAGTTTCTTCCACAACAGGCCACTTACTTTCCCCCGTCACCACCAAGCTCAGAGCCTGGAAGTCCAGATAGACAAGCAGAGATGCTCCAGAATTTAACCCCACCTCCATCCTATGCTGCTACAATTGCTTCTAAACTGGCAATTCACAATCCAAATTTACCCACCACCCTGCCAGTTAACTCACAAAACATCCAACCTGTCAGATACAATAGAAGGAGTAACCCCGATTTGGAGAAACGACGCATCCACTACTGCGATTACCCTGGTTGCACAAAAGTTTATACCAAGTCTTCTCATTTAAAAGCTCACCTGAGGACTCACACTGGTGAAAAGCCATACAAGTGTACCTGGGAAGGCTGCGACTGGAGGTTCGCGCGATCGGATGAGCTGACCCGCCACTACCGGAAGCACACAGGCGCCAAGCCCTTCCAGTGCGGGGTGTGCAACCGCAGCTTCTCGCGCTCTGACCACCTGGCCCTGCATATGAAGAGGCACCAGAACTGA**GCTGCCTTCTGCGGGGCTTGCCTTCTGGCCATGCCCTTCTTCTCTCCCTTGCACCTGTACCTCTTGGTCTTTGAATAAAGCCTGAGTAGGAAGTAGCATAACCCCTTGGGGCCTCTAAACGGGTCTTGAGGGGTTTTTTG**

MYC

**TAATACGACTCACTATAG**GG**AAATAAGAGAGAAAAGAAGAGTAAGAAGAAATATAAGAGCCACC**atgcccctcaacgttagcttcaccaacaggaactatgacctcgactacgactcggtgcagccgtatttctactgcgacgaggaggagaacttctaccagcagcagcagcagagcgagctgcagcccccggcgcccagcgaggatatctggaagaaattcgagctgctgcccaccccgcccctgtcccctagccgccgctccgggctctgctcgccctcctacgttgcggtcacacccttctcccttcggggagacaacgacggcggtggcgggagcttctccacggccgaccagctggagatggtgaccgagctgctgggaggagacatggtgaaccagagtttcatctgcgacccggacgacgagaccttcatcaaaaacatcatcatccaggactgtatgtggagcggcttctcggccgccgccaagctcgtctcagagaagctggcctcctaccaggctgcgcgcaaagacagcggcagcccgaaccccgcccgcggccacagcgtctgctccacctccagcttgtacctgcaggatctgagcgccgccgcctcagagtgcatcgacccctcggtggtcttcccctaccctctcaacgacagcagctcgcccaagtcctgcgcctcgcaagactccagcgccttctctccgtcctcggattctctgctctcctcgacggagtcctccccgcagggcagccccgagcccctggtgctccatgaggagacaccgcccaccaccagcagcgactctgaggaggaacaagaagatgaggaagaaatcgatgttgtttctgtggaaaagaggcaggctcctggcaaaaggtcagagtctggatcaccttctgctggaggccacagcaaacctcctcacagcccactggtcctcaagaggtgccacgtctccacacatcagcacaactacgcagcgcctccctccactcggaaggactatcctgctgccaagagggtcaagttggacagtgtcagagtcctgagacagatcagcaacaaccgaaaatgcaccagccccaggtcctcggacaccgaggagaatgtcaagaggcgaacacacaacgtcttggagcgccagaggaggaacgagctaaaacggagcttttttgccctgcgtgaccagatcccggagttggaaaacaatgaaaaggcccccaaggtagttatccttaaaaaagccacagcatacatcctgtccgtccaagcagaggagcaaaagctcatttctgaagaggacttgttgcggaaacgacgagaacagttgaaacacaaacttgaacagctacggaactcttgtgcgtaa**GCTGCCTTCTGCGGGGCTTGCCTTCTGGCCATGCCCTTCTTCTCTCCCTTGCACCTGTACCTCTTGGTCTTTGAATAAAGCCTGAGTAGGAAGTAGCATAACCCCTTGGGGCCTCTAAACGGGTCTTGAGGGGTTTTTTG**

TFAP2C isoform 1

**TAATACGACTCACTATAG**GG**AAATAAGAGAGAAAAGAAGAGTAAGAAGAAATATAAGAGCCACC**ATGTTGTGGAAAATAACCGATAATGTCAAGTACGAAGAGGACTGCGAGGATCGCCACGACGGGAGCAGCAATGGGAATCCGCGGGTCCCCCACCTCTCCTCCGCCGGGCAGCACCTCTACAGCCCCGCGCCACCCCTCTCCCACACTGGAGTCGCCGAATATCAGCCGCCACCCTACTTTCCCCCTCCCTACCAGCAGCTGGCCTACTCCCAGTCGGCCGACCCCTACTCGCATCTGGGGGAAGCGTACGCCGCCGCCATCAACCCCCTGCACCAGCCGGCGCCCACAGGCAGCCAGCAGCAGGCCTGGCCCGGCCGCCAGAGCCAGGAGGGAGCGGGGCTGCCCTCGCACCACGGGCGCCCGGCCGGCCTACTGCCCCACCTCTCCGGGCTGGAGGCGGGCGCGGTGAGCGCCCGCAGGGATGCCTACCGCCGCTCCGACCTGCTGCTGCCCCACGCACACGCCCTGGATGCCGCGGGCCTGGCCGAGAACCTGGGGCTCCACGACATGCCTCACCAGATGGACGAGGTGCAGAATGTCGACGACCAGCACCTGTTGCTGCACGATCAGACAGTCATTCGCAAAGGTCCCATTTCCATGACCAAGAACCCTCTGAACCTCCCCTGTCAGAAGGAGCTGGTGGGGGCCGTAATGAACCCCACTGAGGTCTTCTGCTCAGTCCCTGGAAGATTGTCGCTCCTCAGCTCTACGTCTAAATACAAAGTGACAGTGGCTGAAGTACAGAGGCGACTGTCCCCACCTGAATGCTTAAATGCCTCGTTACTGGGAGGTGTTCTCAGAAGAGCCAAATCGAAAAATGGAGGCCGGTCCTTGCGGGAGAAGTTGGACAAGATTGGGTTGAATCTTCCGGCCGGGAGGCGGAAAGCCGCTCATGTGACTCTCCTGACATCCTTAGTAGAAGGTGAAGCTGTTCATTTGGCTAGGGACTTTGCCTATGTCTGTGAAGCCGAATTTCCTAGTAAACCAGTGGCAGAATATTTAACCAGACCTCATCTTGGAGGACGAAATGAGATGGCAGCTAGGAAGAACATGCTATTGGCGGCCCAGCAACTGTGTAAAGAATTCACAGAACTTCTCAGCCAAGACCGGACACCCCATGGGACCAGCAGGCTCGCCCCAGTCTTGGAGACGAACATACAGAACTGCTTGTCTCATTTCAGCCTGATTACCCACGGGTTTGGCAGCCAGGCCATCTGTGCCGCGGTGTCTGCCCTGCAGAACTACATCAAAGAAGCCCTGATTGTCATAGACAAATCCTACATGAACCCTGGAGACCAGAGTCCAGCTGATTCTAACAAAACCCTGGAGAAAATGGAGAAACACAGGAAATAA**GCTGCCTTCTGCGGGGCTTGCCTTCTGGCCATGCCCTTCTTCTCTCCCTTGCACCTGTACCTCTTGGTCTTTGAATAAAGCCTGAGTAGGAAGTAGCATAACCCCTTGGGGCCTCTAAACGGGTCTTGAGGGGTTTTTTG**
